# Pubertal timing is a critical dimension of adolescent brain vulnerability

**DOI:** 10.64898/2026.11.02.763578

**Authors:** Hanna Szakács, Ferenc Gombos, Morten L. Kringelbach, Gustavo Deco, Ilona Kovács

## Abstract

Adolescence is a period of rapid brain development, during which biological maturational timing varies substantially between individuals. This variability may drive specific developmental vulnerabilities to adverse experiences with long-term consequences, warranting early identification of at-risk individuals. We utilized ultrasonic bone age measurements to track pubertal maturity, classifying participants by adolescent biological maturity relative to chronological age (accelerated, average, decelerated). Brain hierarchy was quantified as the irreversibility of brain signals in resting state electroencephalography, providing a brain-based method to detect vulnerability patterns. The COVID-19 pandemic presented a unique opportunity to also assess alterations in brain hierarchy following external disruption. We mapped maturity-defined trajectories and rates of hierarchical organization, and their post-pandemic deviation from expected development, capturing two distinct vulnerability forms. First, early maturing adolescents showed persistent reduction in brain hierarchy. Second, those undergoing their fastest phase of development – specifically, decelerated adolescents – showed the strongest disruption in brain hierarchy following the pandemic. These findings suggest that pubertal timing is a critical dimension of individual variation in adolescent brain development and vulnerability. They also emphasize the need for objective maturity measurements to tailor support to individual pubertal timelines, particularly given the secular trend toward earlier puberty and ongoing global ecological crises.

## Introduction

Neuroendocrine changes triggered by puberty fundamentally shape outcomes across cognition^1–3^, mental health^4–6^, behaviour^7–11^, and structural reorganization in the brain^12–15^ during adolescence. The substantial variability of pubertal timing ^3,16^(Fig. 1a) suggests that individuals may experience distinct cerebral developmental paths depending on when pubertal changes first occur. An accelerated maturational timing, characterized by early maturation relative to peers, has been associated with increased risk of psychopathology^4,17–20^, but also with enhanced performance in certain cognitive domains^1,21–23^. Decelerated timing, although less studied, either does not seem to share traits associated with accelerated timing^4,17,19^ or has explicitly shown distinct outcomes in mental health and cognition^1,18,24^, suggesting divergent developmental pathways. Despite extensive literature aiming to evaluate the precise impact of pubertal maturity on cerebral development, these efforts are restricted by critical limitations of current maturity assessments: Tanner staging is subjective and observer-dependent^25,26^ while hormonal measures are volatile due to pulsatile secretion^27^.

**Fig. 1:**
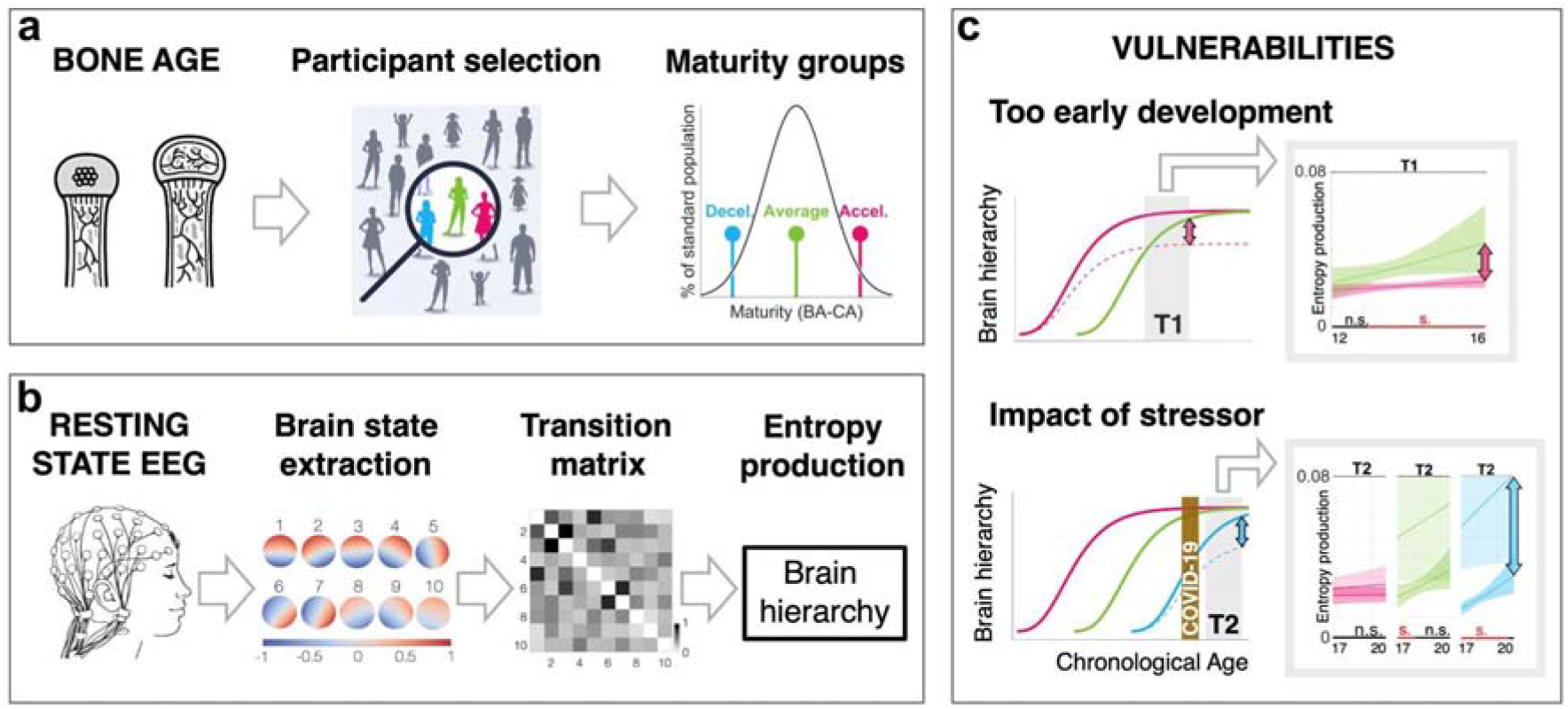
Assessing pubertal timing dependent cortical hierarchy. **a**, After measuring bone age in a large cohort of adolescents, we selected an equal number of participants from three non-overlapping groups of different maturity levels. Bone age is an objective gauge of biological age (BA) that aligns with pubertal hormonal levels. Maturity is defined by the difference between BA and chronological age (CA), and it follows a normal distribution in the standard population. To reliably compare developmental trajectories of adolescents with decelerated (Decel., blue), average (green), and accelerated (Accel., pink) pubertal timing, we balanced the number of participants within each group. **b**, Brain-based assessment of cortical hierarchical development was carried out by recording closed eye resting state EEG. We parsed signals into a sequence of topographically distinct activity patterns, referred to as brain states. The scale denotes cosine similarity values of signals (red: in-phase; blue: counterphase). We calculated entropy production, which is a measure of brain hierarchy, based on the transition matrices between brain states. Brain hierarchy increases with the irreversibility between brain states (darker grey elements in the matrix). **c**, We modelled developmental trajectories of brain hierarchy using the Gompertz function on longitudinal resting state EEG data acquired at two waves (T1: 12-16 yrs.; T2: 17-20 yrs.). First, we asked whether starting puberty earlier (pink lines) leads to a long-term deviation (pink arrow) from the typical trajectory (green). We found that the developmental trajectory of accelerated adolescents deviates significantly from mid-adolescence on in T1 (color shaded areas are confidence intervals; n.s. denotes non-significant, s. (red) denotes significant deviations as a function of CA). Second, we asked whether the difference in the steepness of developmental curves at the time of COVID-19 scales with the deviation from the expected developmental trajectory at T2 (blue arrow). We used pre-pandemic trajectories to predict expected development in T2 (lighter color shading) and found that the late developing group (blue lines) with the steepest curve diverged to the greatest extent.

To resolve uncertainty around the exact relationship between maturational timing and development, we introduced ultrasonic bone age measurements^22^ as a proxy for biological age (BA) in adolescent research (Fig. 1a). Skeletal development closely aligns with pubertal hormone levels^28,29^, providing an objective and stable assessment of biological age in adolescence. In this paradigm, we define maturity as the difference between chronological age (CA) and BA, enabling the classification of participants into accelerated, average, and decelerated maturity groups. Relying on this grouping approach, we previously demonstrated maturity-related effects in cognition^22,30^, motor function^31^, cortical hierarchization^32^, and emotion regulation^33^.

However, recent neuroimaging studies leveraging longitudinal trajectories highlight that comparison of group-level, aggregate outcomes across groups may obscure dynamic changes in brain structure^34–37^ and function^35,38,39^ or periods of heightened susceptibility to behavioural and psychological problems during development^40,41^, all of which are detectable only when analyzed over extended time spans. Conversely, outlining developmental trajectories across subpopulations over a wide age range reveals dynamic developmental features specific to these groups^42,43^. In this work, we rely on developmental trajectory modelling to uncover distinct outcomes in brain hierarchy and dynamic characteristics such as the rate of development within an assessed time window (referred to here as maturity-related momentum). We employ the Gompertz function, a 3-parameter nonlinear function that provides an ideal solution for the longitudinal modelling of complex developmental trajectories in the brain^44–49^.

Here, we trace brain development through the changing level of cortical hierarchization in resting state EEG recordings (Fig. 1b). The level of functional hierarchy in the human brain has been proposed to be indexed by the temporal asymmetry or irreversibility in the flow of neural events, a method grounded in the thermodynamic arrow of time^50^. This concept denotes that the reverse of the observed processes of a system is statistically highly unlikely to occur^51^. In the brain, irreversibility may manifest in multiple ways, such as asymmetric information flow between brain regions^52^, the asymmetric probability of transitions in neural state space^53^, or the similarity of forward and reverse brain signal time series^54^.

Increased hierarchy in the brain has been observed during cognitive and physical task performance compared to rest^52,53,55^. On the other hand, decreased hierarchy has been observed in disorders of consciousness^56^, attention-deficit hyperactivity disorder and bipolar disorder^52^, Parkinson’s and Alzheimer’s disease^57,58^, and epilepsy^58,59^. In a developmental sample of school-aged children, resting-state irreversibility increased with age in students enrolled in a Montessori school, coinciding with significantly higher cognitive performance relative to traditionally schooled peers^60^. In sum, irreversibility – whether measured during task performance or at rest – is considered to be an indicator of the efficiency and integrity of brain function orchestration^61^.

In this study, we quantify irreversibility during resting state by parsing each EEG recording into topographically distinct activity patterns referred to as brain states (Fig. 1b). These recurring patterns of synchronized activity represent the instantaneous global phase relationships of EEG signals across the cortex^62^. We assess the extent to which forward and reverse transitions between EEG-defined brain states follow a directional pattern rather than occurring symmetrically. The level of directional asymmetry is quantified via entropy production calculations, indexing hierarchical brain organization^53,63^.

In previous work, our group evaluated resting state EEG-derived brain hierarchy in a cohort of 12-13-year-old adolescent females^32^, sorted into accelerated, average, and decelerated groups according to maturity assessed in adolescence. Comparisons between maturity groups showed that an average maturational timing was associated with a significantly higher level of entropy production, and both accelerated and decelerated groups showed lower levels of cortical hierarchy. We termed this finding the Goldilocks rule of brain development, reflecting that brain hierarchical organization in adolescence seems to favour a maturational timing that is just right. However, this interpretation may imply that similar levels of hierarchical organization across atypical groups are generated by the same underlying neurodevelopmental process.

Here, we propose that accelerated and decelerated maturational timing might reflect fundamentally distinct neurodevelopmental paths. Specifically, accelerated maturational timing might be associated with persistent reduction in hierarchical organization, while decelerated timing might involve a transient period of reduced hierarchy that recovers over time. Additionally, periods of rapid maturity-related momentum may signal increased vulnerability to psychological distress (Fig. 1c).

The COVID-19 pandemic represents the most recent large-scale disruption that exposed an entire generation of adolescents to multiple external stressors: lockdown measures, including school closures^64^, disrupted their daily lives, taking a toll on their mental health: symptoms of depression and anxiety increased^65–68^, while reported irritability and anger also surged^69^. Youth also experienced a pandemic-related decline in their educational gains^70^ and cognitive performance, both in domain-specific areas^71^ and in terms of overall intelligence measures^72^. The far-reaching consequences of the pandemic are also reflected in the interruption of normative adolescent brain development, with a decreased activation in response to affective stimuli in the prefrontal cortex^73^, and accelerated maturation in multiple brain regions including the hippocampus^71,74,75^, amygdala^71,74^, and prefrontal cortex^73,75^, and across widespread cortical regions^76^. These reports on increased cortical developmental rates are especially alarming as accelerated biological aging is a well-documented aftermath of adversity^77–80^. In sum, the evidence reflects detrimental effects linked to the pandemic across a broad range of domains in adolescent development. However, whether maturational timing itself shapes differential vulnerability to the prolonged stress of the pandemic and related preventive measures, remains unknown.

In this work, we examine two hypothesized, maturity-related vulnerabilities in hierarchical cortical development, relying on a longitudinal cohort of female participants. First, we assess whether accelerated timing is associated with persistently reduced hierarchy, and second, we gauge whether maturational timing and maturity-related momentum modify the extent of deviation from expected brain hierarchy development following large-scale external stress.

## Results

We assembled an all-female longitudinal cohort with two data acquisition waves: first in adolescence (T1; 11.60–15.97 years), before pandemic-related school closures occurred, and again approximately 5 years later, in emerging adulthood (T2; 16.91–19.99 years), after pandemic restrictions were lifted^81^. At both data acquisition waves, we collected resting state EEG recordings from which we calculated entropy production, repeating our previously published procedure^32^. We estimated trajectories of hierarchical organization by modelling age-related patterns of entropy production in each maturity group within each wave. To this end, we used the Gompertz function, with uncertainty incorporated through bootstrapping (see section *Gompertz function fitting*, and equations 3 and 4 in *Methods*). In each analysis, trajectories were statistically compared pairwise using a windowed area integration method (area-under-the-curve; AUC; see section *Statistical comparison of Gompertz functions* and equation 5 in *Methods*) to assess temporally resolved differences in neurodevelopmental change over time.

### Accelerated maturity is associated with persistent reduction in hierarchical brain organization

We first compared trajectories pairwise across maturity groups at T1 to assess maturational timing-specific differences in developmental patterns across the adolescent age range. We assigned T1 participants to maturity groups based on maturational timing, as specified in Fig. 1a, creating decelerated (BA<CA-0.5, mean BA-CA: −1.01, s.d.=0.48; N=31), average (CA-0.5≤BA≤CA+0.5, mean BA-CA: −0.01, s.d.=0.28; N=28), and accelerated (BA>CA+0.5, mean BA-CA: 0.94, s.d.=0.36; N=28) maturity groups. We then conducted pairwise comparisons of maturity group-specific developmental trajectories, evaluating AUC over 0.25-year-wide CA windows between 11.60 and 15.75 years, and a 0.21-year-wide window between 15.76–15.97 years.

Comparison of developmental trajectories of the average and accelerated maturity groups (Fig. 2a) did not yield significant results across windows spanning from 11.60 to 12.63 years; however, the trajectories showed significant divergence across windows spanning from 12.64 to 15.97 years, with the average group expressing higher entropy production values. Mean AUC values and 95% CIs of assessed CA windows can be found in Table S1.

**Fig. 2:**
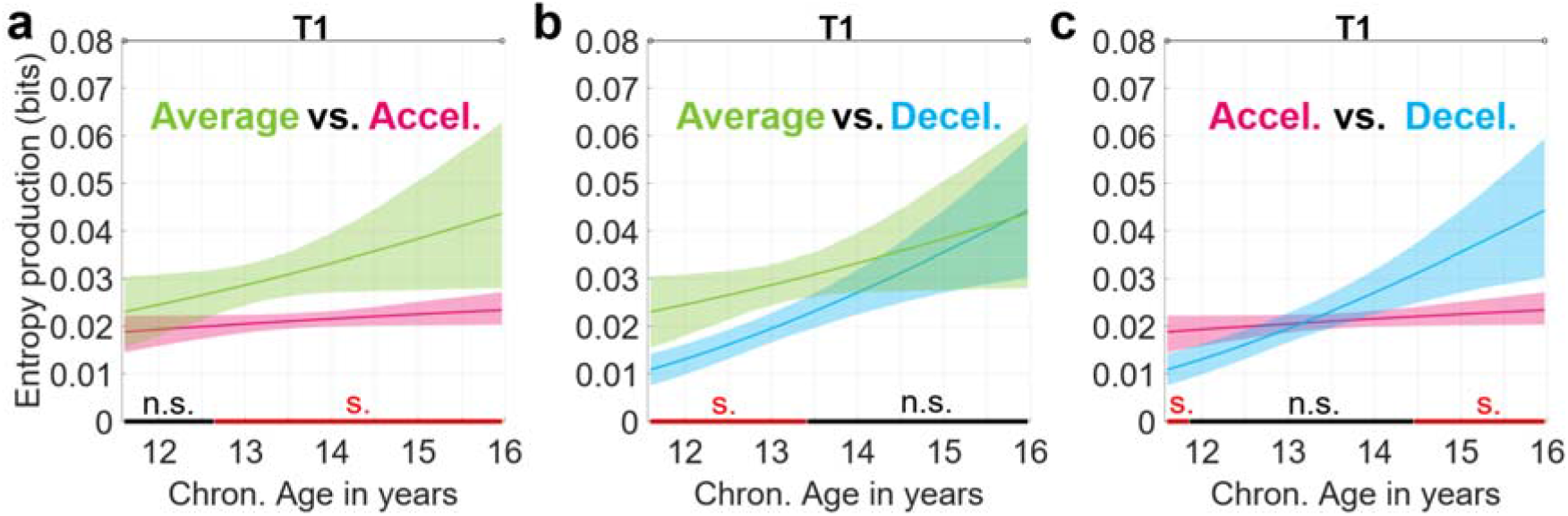
Pairwise comparison of developmental trajectories across T1 maturity groups. Each panel shows pairwise comparisons of maturity group-specific developmental trajectories of entropy production (y-axis) over chronological age (x-axis) at T1, color-coded according to maturity group: average (green), accelerated (pink), and decelerated (blue). Trajectories were estimated by fitting Gompertz functions to 2000 bootstrap resamples within each maturity group. Mean Gompertz functions (lines) and 83.5% CIs (shading) were calculated from bootstrap-derived fits. Trajectories were compared via windowed area integration. Red (s.) marks significant windows; black (n.s.) marks non-significant windows at *p*<0.05. **a**, Average vs. accelerated trajectories showed significant divergence across windows spanning from 12.64 to 15.97 years, with the average group displaying higher entropy production. **b**, Average vs. decelerated trajectories showed significant divergence across windows spanning from 11.60 to 13.41 years, with the average group displaying higher entropy production. Then, trajectories converged over windows starting at 13.42 years, after which comparisons were non-significant. **c**, Accelerated vs. decelerated trajectory comparisons revealed a developmental shift. The trajectory of the accelerated group displayed significantly higher entropy production across windows between 11.60 and 11.85 years. Trajectories converged over windows spanning from 11.86 to 14.45 years, then diverged significantly again across windows covering the interval between 14.46 and 15.97 years as the decelerated group surpassed the accelerated group.

Next, we compared the developmental trajectories of the average and decelerated maturity groups (Fig. 2b). This comparison revealed a significant difference over windows spanning from 11.60 to 13.41 years, with the average group exhibiting higher entropy production values. Then, trajectories converged and did not show significant differences across windows between 13.42 and 15.97 years. Mean AUC values and 95% CIs can be found in Table S2.

Comparison of developmental trajectories (Fig. 2c) of the decelerated and accelerated groups revealed an intriguing shift over time. Initially, trajectories diverged significantly in the window spanning from 11.60 to 11.85 years, with the accelerated group demonstrating higher entropy production values. Following this brief divergence, the trajectories converged across windows spanning from 11.86 to 14.45 years. Then, trajectories again diverged significantly over windows from 14.46 to 15.97 years; however, the decelerated group demonstrated higher entropy production values over this period, surpassing the accelerated group and reversing the initial pattern in terms of hierarchical brain organization. Mean AUC values and 95% CIs are presented in Table S3.

Following the pairwise comparison of developmental trajectories, we assessed whether the pace of developmental change (momentum) is influenced by adolescent maturational timing. We first calculated the mean slope of each trajectory to describe overall momentum (decelerated mean slope: 0.0076 bits/year, 95% CI: 0.0027–0.0137; average mean slope: 0.0047 bits/year, 95% CI: 0.0000–0.01258; accelerated mean slope: 0.0011 bits/year, 95% CI: 0.0000–0.0034). Then, to formally assess differences in maturity-related momentum, we compared slopes across the same CA windows as defined in trajectory comparisons (see section *Slope calculation* in *Methods*).

The pairwise windowed comparison of slopes revealed that the decelerated group is characterized by a significantly steeper slope over the entire T1 CA range compared to the accelerated group. The significant difference in slopes persisted even during periods when developmental trajectories converged, revealing fundamentally distinct patterns of maturity-related momentum between the atypical maturity groups. In contrast, the comparison of slopes between the accelerated vs. average, and decelerated vs. average groups did not show significant divergence within any of the predefined CA windows. Window-specific slope values and 95% CIs are presented in Tables S4, S5 and S6.

To assess the robustness of our results to window width, we performed sensitivity analyses using 0.50-year, 1.0-year, and 1.5-year windows for both modelled trajectory and slope comparisons across the adolescent age range. Sensitivity analyses reproduced the patterns reported in the main text. AUC and slope values, accompanied by 95% CI values and an indication of significant windows, are reported in Tables S15-S20.

These results support our hypothesis that atypical maturity groups are linked to distinct developmental patterns: the initial reduction in entropy production characterising the decelerated group compared to typically maturing peers was transient and disappeared over time, driven by rapid maturity-related momentum, while the deviation observed in the accelerated group persisted across adolescence.

### Deviation in hierarchical brain organization following the pandemic

To assess alterations in cortical hierarchical organization following the pandemic, we projected the pre-pandemic T1 trajectory into the T2 age range to model a normative reference trajectory reflecting nondisrupted development. Then, we compared the reference trajectory to the observed, post-pandemic T2 trajectory to quantify developmental deviation. We evaluated the divergence of trajectories over 0.25-year-wide CA windows between 16.91 and 19.76 years and a 0.22-year-wide CA window between 19.77 and 19.99 years. At T2, participants retained their T1 maturity group classifications (decelerated: N=19, mean CA: 18.53, s.d.=0.78; average: N=18, mean CA: 18.06, s.d.=0.62; accelerated: N=19, mean CA: 18.12, s.d.=0.86). Participants of both samples are represented in Fig. 3a as color-coded data points, denoting CA and dates of EEG data acquisition. Yellow shading represents school closing measures in Hungary at three severity levels, reported daily between 1/1/2020 and 12/31/2021^64^.

**Fig. 3:**
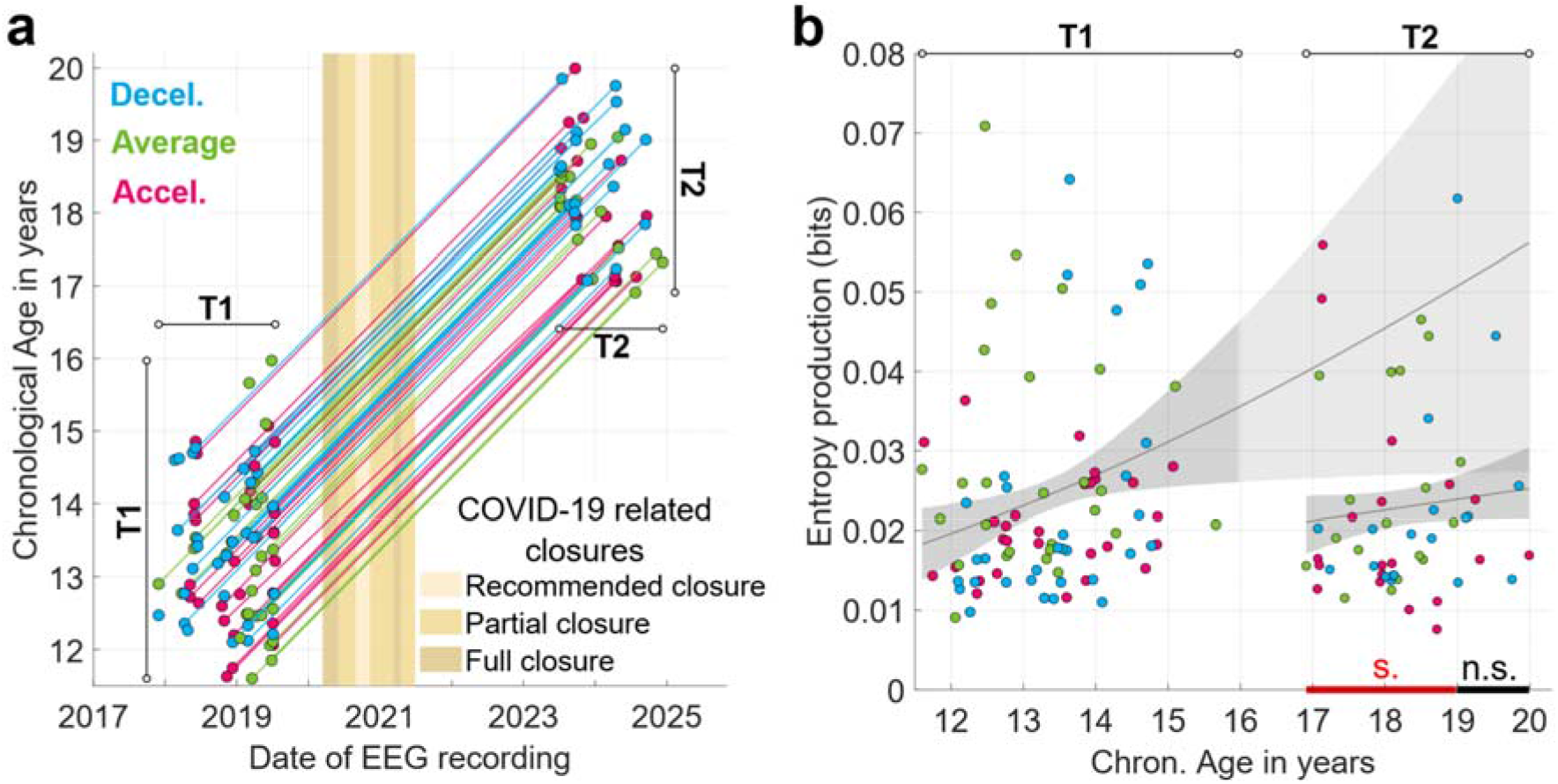
Hierarchical brain organization falls below expected levels in post-pandemic assessments. **a**, Data acquisition dates (x-axis) and chronological age (y-axis) of participants, color-coded by maturity group: decelerated (blue), average (green), and accelerated (pink). Participants were assessed in two waves: once in adolescence, before the pandemic (T1; left side of panel) and again in emerging adulthood (T2; right side of panel), after COVID-19 restrictions were lifted (yellow shading). Maturity group assignment was maintained from T1 to T2. Straight lines connect participants with data collected in both waves. **b**, Comparison of T1-based normative developmental trajectory versus observed T2 trajectory for entropy production (y-axis) across chronological age (x-axis). Horizontal lines at top delineate T1 and T2 age ranges. Dark grey lines and shading show mean Gompertz functions and 83.5% CIs of each sample. The T1 trajectory was projected into the T2 age range (lighter shading) to model undisrupted development. AUC analysis compared projected and observed trajectories over the T2 age range, revealing significant divergence at *p*<0.05 (red line; s.) across windows spanning from 16.91 to 18.98 years, with T1 projections exceeding T2 observations, indicating a disruption in brain hierarchy in post-pandemic assessments. No significant divergence occurred within windows between 18.99–19.99 years (black line; n.s.).

Post-pandemic observations fell significantly below normative projections within windows spanning from 16.91 to 18.98 years (Fig. 3b), signifying a setback in entropy production following the COVID-19 pandemic and related school closures. The trajectories then converged in windows covering the age range between 18.99 and 19.99 years. This result indicates a disruption in the normative developmental process of cortical hierarchical organization between the pre- and post-pandemic periods. Table S7 reports the mean AUC values and corresponding 95% CIs for each assessed CA window.

We performed sensitivity analyses using 0.50–year, 1.0–year, and 1.5–year windows to assess whether the comparison results of normative projections and post-pandemic measurements are robust to the windowing approach. Sensitivity analyses confirmed patterns reported here. AUC and 95% CI values, with an indication of significant windows, are reported in Table S21.

While this result shows an overall deviation in cortical hierarchical organization in post-pandemic assessments, it does not resolve whether the alteration is uniform across variations of maturational timing. To assess maturational timing-specific patterns, we first compared maturity group means within T1 and T2 samples to characterize group-level differences in entropy production across adolescence and emerging adulthood.

### Vulnerability scales with maturity-related momentum in the adolescent brain

To compare mean levels of entropy production across maturity groups within T1 and T2 samples, we calculated group-level average entropy production using inverse-variance weighted averaging (equations 7 and 8 in *Methods*) and assessed statistical differences via two-sample z-tests (equations 9 and 10 in *Methods*).

At T1, the average maturity group (0.0240 bit; 95% CI: 0.0213–0.0267) showed significantly higher group-level entropy production compared to both the decelerated group (0.0181 bit; 95% CI: 0.0158–0.0204; *p*=0.001; Hedges’ *g*: 0.85, large effect) and the accelerated group (0.0194 bit; 95% CI: 0.0170–0.0219; *p*=0.015; Hedges’ *g*: 0.65, medium effect). These results replicate our earlier findings^32^, reported on a sample of 12- and 13-year-old adolescent female participants. Inverse-variance weighted averages, 95% CIs and statistically significant comparisons in the T1 sample are depicted in bar charts in Fig. 4 (top row), alongside maturity group-specific network graphs showing brain state transition dynamics (see figure caption for details). Additional descriptive data on T1 groups are reported in Tables S8 and S9.

**Fig. 4:**
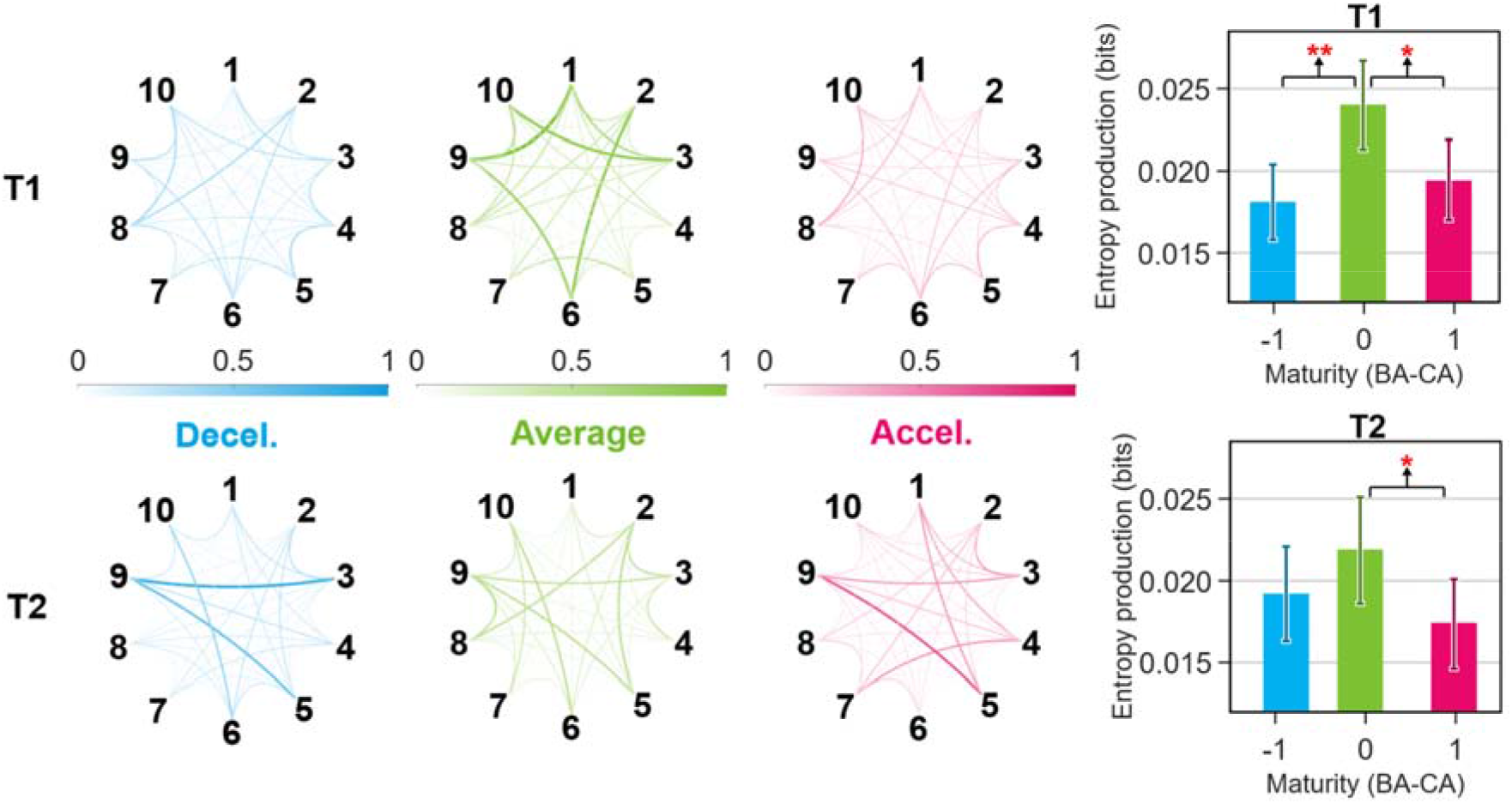
Comparison of group-level hierarchical brain organization within samples. **Top row**: T1 (adolescence, pre-pandemic); **Bottom row**: T2 (emerging adulthood, post-pandemic). On the left: graph representations of brain state transition dynamics contributing to entropy production, color-coded according to maturity group. Nodes represent the 10 brain states derived from resting state EEG (extracted separately for T1 and T2), and edges represent the contribution of pairwise brain state dynamics to total entropy production. Edge thickness and darkness indicate contribution to entropy production (normalized 0-1 scale across all graphs). Right panels: inverse-variance weighted average entropy production by maturity group (x-axis; bars: average; error bars: 95% CI; asterisks: significant differences from two-sample z-tests). At T1, the average group (0.0240 bit; 95% CI: 0.0213–0.0267) showed significantly higher entropy production (y-axis) than both decelerated (0.0181 bit; 95% CI: 0.0158–0.0204; *p*=0.001, *g*=0.85) and accelerated (0.0194 bit; 95% CI: 0.0170–0.0219; *p*=0.015, *g*=0.65) groups. At T2, the significant difference between the average (0.0219 bit; 95% CI: 0.0186–0.0251) and accelerated (0.0174 bit; 95% CI: 0.0146–0.0201) groups persisted (*p*=0.038, *g*=0.69). Pairwise comparisons of random-effects weighted means, and Mann-Whitney U comparisons corroborated these findings at T1 but not at T2.

To complement comparisons based on inverse-variance weighted averages, we also calculated group means using random-effects weighting^82^ and without applying a weighting method, then compared groups using two-sample z-tests and Mann-Whitney U tests, respectively. Comparison of random-effects weighted means revealed significantly higher group-level entropy production in the average maturity group (0.0278 bit; 95% CI: 0.0228– 0.0328) compared to the decelerated (0.0202 bit; 95% CI: 0.0165–0.0239; *p*=0.017; Hedges’ *g*: 0.47, small effect) and the accelerated (0.0194 bit; 95% CI: 0.0170–0.0219; *p*=0.003; Hedges’ *g*: 0.63, medium effect) groups. Unweighted arithmetic means and Mann-Whitney U tests corroborated this pattern, as the average maturity group (0.0305 bit; 95% CI: 0.0236– 0.0374) showed significantly higher levels of entropy production than the decelerated (0.0232 bit; 95% CI: 0.0178–0.0286; *U*=279.000, *p*=0.019, *r*=0.31, medium effect) and the accelerated (0.0209 bit, 95% CI: 0.0184–0.0234; U =265.000, p=0.037, r=0.28, small effect) groups. In sum, all three comparisons reflect significantly higher entropy production in the average maturity group at T1.

At T2, comparison of the inverse-variance weighted average entropy production of the average maturity group (0.0219 bit; 95% CI: 0.0186–0.0251) and the accelerated maturity group (0.0174 bit; 95% CI: 0.0146–0.0201) remained statistically significant (*p*=0.038, Hedges’ *g*: 0.69; medium effect), with the accelerated group showing lower group-level entropy production. Comparison of the T2 average and decelerated groups (0.0192 bit; 95% CI: 0.0163–0.0221) was not significant. Inverse-variance weighted averages, 95% CIs and statistically significant comparisons in the T2 sample are depicted in bar charts on the bottom row of Fig. 4, accompanied by maturity group-specific network graphs illustrating brain state transition dynamics. Additional descriptive data on T2 groups are shown in Tables S10 and S11. Random-effects weighted mean comparisons and Mann-Whitney U tests at T2 did not indicate significant differences between maturity groups. However, random-effects weighted means and unweighted arithmetic means reflected a trend of higher entropy production in the average group and a relative reduction in the atypical groups at T2. Means, 95% CIs, *U* and *p* values, and effect sizes for both T1 and T2 are reported in Fig. S1 in the Supplementary material.

Overall, when assessed at the group-level, both atypical groups showed significantly lower entropy production compared to the average group at T1. At T2, the trend of these differences persisted but were not statistically significant.

However, comparisons of group means obscure long-term, time-resolved developmental dynamics. Our trajectory-based analyses showed that decelerated adolescents exhibited rapid maturity-related momentum with steep increases in entropy production across adolescence, whereas accelerated adolescents showed attenuated momentum and a flat trajectory, leading the decelerated group to surpass the accelerated group in cortical hierarchical organization (Fig. 2c).

To track dynamic, time-resolved developmental changes, we compared T1-based normative projections with observed T2 trajectories within each maturity group, assessing the extent of deviation from expected developmental trajectories of cortical hierarchical organization across maturational timing. We conducted AUC analyses over 0.25-year-wide CA windows between 16.91 and 19.76 years, and a 0.22-year-wide CA window between 19.77 and 19.99 years within each maturity group. Results of this analysis are depicted in Fig. 5.

**Fig. 5:**
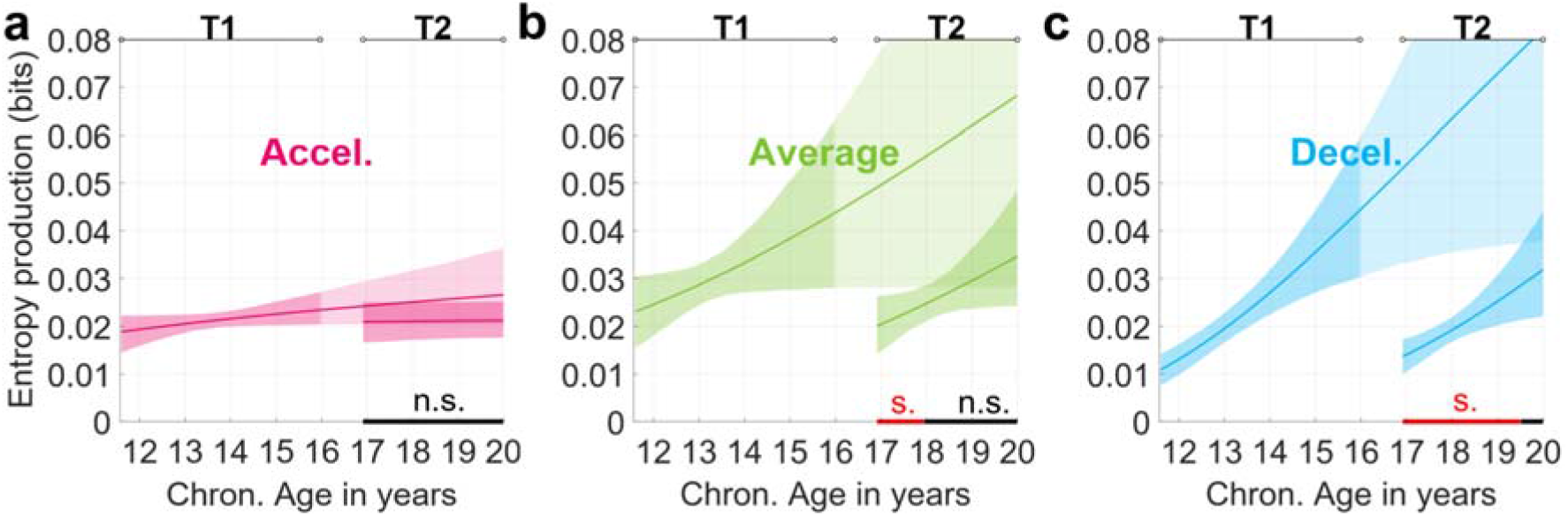
Projected and observed developmental trajectories reveal maturity group-specific vulnerability and maturity-related momentum. Panels show comparisons of T1-based projected trajectories versus observed T2 trajectories for each maturity group. X-axis: global CA range; y-axis: entropy production. Color-coded lines show mean Gompertz functions of T1 and T2 samples, shading represents 83.5% CIs. The T1 and T2 CA ranges are marked at the top of each panel. T1 trajectories are projected (lighter shading) into the T2 age range to model expected development. AUC analysis compared projected and observed trajectories. Red markers (s.) indicate windows with significant divergence; black markers (n.s.) indicate non-diverging windows at *p*<0.05. **a**, The T1-based projection and observed T2 trajectory of the accelerated group do not show significant divergence, illustrating that an accelerated maturational timing is associated with a persistent deviation in hierarchical brain organization. **b**, The T1-based projection and observed T2 trajectory of the average group shows significant divergence within windows spanning from 16.91 to 17.94 years, with T2 observations below T1 projections, indicating considerable deviation from expected development. **c**, The comparison of the T1-based projection and observed T2 trajectory of the decelerated group showed the largest number of windows over which a significant divergence was detected (16.91-19.50 years), indicating the most extensive alteration in hierarchical brain organization in post-pandemic assessments, suggesting that vulnerability coincides with rapid neurodevelopment.

We first compared the projected and observed developmental trajectories within the accelerated maturity group (Fig. 5a). The divergence from the expected T1-based trajectory is not significant within any of the examined CA windows, indicating that the hierarchical cortical organization of the accelerated group did not diverge from its expected trajectory following the pandemic. Table S12 reports mean AUC values and 95% CIs for each assessed window.

In the next analysis, we compared the T1-derived normative developmental trajectory and the observed T2 trajectory of the average maturity group over the T2 age range (Fig. 5b), revealing a significant divergence in the windows spanning from 16.91 to 17.94 years, with the normative trajectory predicting significantly higher entropy production values over this time period. Comparison of the T1-based projection and the observed T2 trajectory did not yield significant results in subsequent windows, starting from 17.95 years. Mean AUC values and 95% CIs for each assessed window are reported in Table S13.

Lastly, comparison of the T1-based projected developmental trajectory and observed T2 trajectory within the decelerated group (Fig. 5c) showed a significant divergence across windows spanning from 16.91 to 19.50 years, with the observed T2 trajectory persistently lagging behind the T1 projection over this interval. This comparison did not yield significant results in windows beyond 19.50 years. This result reflects the most pronounced deviation from expected development in post-pandemic assessments among the maturity groups. Mean AUC values and 95% CIs for each assessed window are reported in Table S14.

We additionally tested the robustness of these comparisons to window width by repeating the analyses with 0.50-year, 1.0-year, and 1.5-year windows. Results were consistent with those reported above. AUC and 95% CI values, along with significance indicators, are reported in Tables S22-S24.

Overall, our results reveal that the vulnerability of developmental trajectories is modulated by adolescent maturational timing, specifically, by maturity-related momentum. The accelerated group demonstrated a flat developmental trajectory at T1 and did not show significant divergence from its projected trajectory in terms of hierarchical brain organization following the pandemic. In stark contrast, the decelerated group exhibited rapid maturity-related momentum at T1, indicating ongoing hierarchical organization that proved highly susceptible to disruption. These findings indicate that rapid maturity-related momentum reflects both fast-paced neurodevelopment and heightened susceptibility to disruptive events.

## Discussion

In the present study, we utilized an objective, brain-based method to examine how variations across adolescent maturational timing drive the course of cortical hierarchical organization, and to detect vulnerabilities tied to these variations. We assessed the maturational timing-specific developmental trajectory and maturity-related momentum of cortical hierarchical organization utilizing a longitudinal female sample to address these questions. We identified two distinct forms of vulnerability linked to maturational timing. First, accelerated maturational timing is associated with a persistent reduction in cortical hierarchical organization. Second, susceptibility to disruption in developmental trajectories aligns with periods of peak maturity-related momentum – denoting the rate of development within an assessed time window –, which in this cohort was most pronounced in the decelerated group in pre-pandemic assessments.

We first assessed how maturational timing shapes the trajectory of cortical hierarchization in adolescence. To this end, we estimated neurodevelopmental trajectories by modelling age-related patterns of entropy production within each maturity group (accelerated, average, decelerated). We then compared trajectories pairwise across maturity groups. Despite similar initial levels, the average maturity group significantly surpassed accelerated adolescents in terms of entropy production. The trajectory of the decelerated group lagged significantly behind that of the average group at first, but then trajectories converged, erasing the initial gap of the decelerated group relative to typically maturing peers, reflecting a transient period of reduced hierarchy that recovered through rapid maturity-related momentum. Strikingly, the comparison of accelerated and decelerated developmental trajectories revealed markedly different developmental courses: the accelerated group first exceeded the decelerated group in terms of cortical hierarchical organization, however, this pattern was reversed as the decelerated group – characterized by rapid maturity-related momentum – surpassed the accelerated group during adolescence. In contrast, consistently low entropy production values in combination with a flat maturity-related momentum within the accelerated group reflected a persistent reduction in entropy production, indicating an enduring trajectory shift toward lower brain hierarchy within this group.

These findings provide evidence that the similar group-level differences in atypical adolescent maturity groups compared to typically maturing peers observed in our previous work^32^, and replicated here, are associated with distinct underlying developmental processes. Specifically, spontaneous resting state activity in the decelerated group indicated rapid neurodevelopment, while the accelerated group exhibited a stable, persistently low-hierarchy state over adolescence. In sum, the accelerated maturity group seems to be characterized by a persistent trajectory-level alteration in hierarchical brain organization, while the decelerated group appears to express a transient delay in its developmental trajectory.

Reduced irreversibility during resting state was identified in individuals diagnosed with epilepsy^58,59^, Alzheimer’s disease^57^, Parkinson’s disease^58^, and schizophrenia^58,83^ compared to healthy controls, indicating that temporal asymmetry in neural events of spontaneous brain dynamics is a marker of optimal brain functioning, whereas pathology is characterized by a reduction in irreversibility. In the present work, a persistently low-hierarchy resting state activity associated with accelerated maturational timing might index inefficient functioning or even risk for pathology^58^. Indeed, previous studies linked accelerated maturational timing relative to peers to a wide range of internalizing problems such as depressive symptoms^18,19,84–86^, eating disorders^20^, anxiety^84^, and behavioural difficulties such as self-reported criminality, substance use problems, social isolation, and early sexual behaviour^84^ in adolescence. Additionally, a few studies confirmed the persistence of early puberty-related depressive symptoms^84,87^ and antisocial behaviour^87^ into young adulthood.

Puberty-dependent cerebral remodelling^13–15,19,48,88,89^, including reductions in cortical thickness^19,88^ and synaptic density^48,89^, suggests that early maturation might alter normative developmental trajectories of these processes, potentially leading to sustained suboptimal functioning. For example, an early pubertal timing has been linked to advanced cortical thinning in the dorsal attention, default mode, frontoparietal and posterior multimodal networks^88^ and within temporal, frontal and parietal regions^19^, possibly disrupting normative development and adaptability of regions involved in higher-order cognition and emotional processing.

Following the comparison of adolescent trajectories, we assessed the extent to which trajectories of hierarchical brain organization diverged from their projected course following external disruption, taking advantage of the unique occurrence of a global pandemic. We extrapolated pre-pandemic developmental trajectories to serve as normative references, then compared them to post-pandemic observations to quantify deviation from nondisrupted development following the COVID-19 pandemic. Our results showed significant disruptions in the normative advancement of cortical hierarchical organization in post-pandemic measurements. These results align with the mounting evidence of pandemic-related interruptions to typical cerebral development^71,73–76^. In terms of alterations in brain architecture, multiple studies showed that the pandemic accelerated maturation in multiple brain regions including the hippocampus^71,74,75^, amygdala^71,74^ and prefrontal cortex^73,75^, a phenomenon consistent with accelerated biological aging as a consequence of adversity^77–80^. In addition to findings related to anatomical changes, the pandemic was also linked to heightened psychological distress among adolescents, reflected in increases in anxiety, depression, irritability and anger^65–69^. Besides psychological well-being, cognitive performance also showed pandemic-related reductions^70–72^, suggesting that this large-scale environmental disruption is linked to developmental alterations in a broad set of domains among adolescents of this generation.

Critically, trajectory-level deviations from expected development in post-pandemic assessments were not uniform among maturity groups in our sample, as maturational timing modulated their magnitude. Susceptibility to developmental disruption coincided with maturity-related momentum: the developmental trajectory associated with accelerated timing showed attenuated maturity-related momentum at the time pandemic-related preventive measures began^64^ and did not show significant divergence from its expected course, whereas those with decelerated timing, characterized by rapid neurodevelopmental change before the pandemic, exhibited substantial deviation from their projected developmental course. Importantly, the reliable detection of maturity-driven vulnerability patterns was contingent upon an adolescent sample screened for maturational timing before the pandemic, enabling classification by maturational timing that is unconfounded by this external disruption.

Several studies frame adolescence as a sensitive period for the development of heteromodal association cortices^90–93^ as these regions generally experience a protracted period of neuroplasticity, a developmental feature supporting prolonged experience-dependent refinement but also marking a period of heightened vulnerability^94,95^. Specifically, association cortices governing mentalizing and socioemotional functions – e.g., posterior cingulate cortex, temporo-parietal junction, and medial prefrontal cortex – undergo rapid neural reorganization during adolescence^96,97^, potentially rendering them particularly susceptible to environmental disruption, such as pandemic-related stress and isolation from peers. Critically, periods of increased associative cortical neuroplasticity in adolescence are suspected to be strongly influenced by the timing of puberty^95^, consistent with the maturational timing-specific vulnerabilities observed in our study. Within this framework, rapid maturity-related momentum characterising the decelerated group before the pandemic may index an open sensitive period, consistent with the greater deviation from expected developmental trajectories observed in our sample in post-pandemic assessments.

Our study has a few notable limitations. First, it is characterized by a 36% attrition rate, resulting in a lower number of participants in the second wave of data acquisition. However, the achieved retention is comparable to other adolescent cohorts with large intervals between data collection waves^98,99^. Relatedly, the group sizes (N=87 at T1, N=56 at T2; corresponding to 28–31 and 18–19 participants per maturity group, respectively) limit the precision of the modelled trajectories. Importantly, bone age assessment and longitudinal EEG recording are labor-intensive and constrain cohort size, but provide objective, individual-level indices of maturational timing and hierarchical brain organization. Second, we included female participants only, a deliberate choice made to avoid confounding effects stemming from delayed puberty onset in males^100,101^ and from sex-specific developmental patterns of brain morphology^102,103^. Additionally, we selected participants from similar socioeconomic backgrounds, which limits the generalizability of our findings to socioeconomically diverse populations. Importantly, this variable was controlled for as it has been associated with alterations in the pace of brain development^104,105^. Third, trajectories of hierarchical brain organization were modelled at the group level rather than individually. As such, between-person variability in hierarchical brain organization may have contributed to the trajectory shapes. Crucially, the COVID-19 pandemic prevented the repeated assessment of our participants; the group-level modelling approach was therefore adopted to maximize the sample size and age range over which developmental trajectories could be defined. Fourth, as bone age was assessed only once during adolescence, within-individual variation in this metric might have introduced random noise into our dataset. Nevertheless, the single-timepoint classification was sufficient to reveal maturity-specific patterns in both the development and vulnerability of cortical hierarchical organization.

Several important directions for future research emerge from our findings. First, characterizing how entropy production relates to cognitive and behavioural measures across variations of adolescent maturational timing represents an important next step. Prior work established the connection between irreversibility and cognitive function^52,53,55^, yet whether these findings replicate in an adolescent cohort, especially one where maturational timing is identified, remains unknown. Second, the characterization of individual developmental trajectories in addition to group-level trajectories could potentially reveal fine-grained differences between individuals across the maturity spectrum. Third, recent findings show that social isolation, even for short periods and under controlled circumstances, alters core components of reward processing^106^ and increases threat learning^107^ in adolescents, raising the question of how prolonged, real-life isolation may influence vulnerability to reward-related risk-seeking and to threat perception-related disorders (e.g., anxiety and phobias) as a function of maturational timing. In general, future research examining developmental variations should account for the time-resolved dynamics of maturational timing-specific development, as apparent similarities in outcomes may mask fundamentally different underlying neurodevelopmental processes.

Recent decades have seen a rise in reported mental health conditions and neurodevelopmental diagnoses among adolescents^108–114^. It is unclear whether this trend reflects improved recognition and diagnostic practices^108,114^ or a genuine increase in prevalence due to various factors, e.g. digital media use^115,116^, academic pressure^108,115,117^ or lifestyle^108,115^. Critically, these studies do not consider the role of maturational status in these outcomes. This oversight is particularly important given that pubertal neuroendocrine processes heavily influence mental health^4–6^, cognition^1–3^ and behaviour^7–11^. Another secular trend, involving earlier pubertal onset^118,119^, further complicates this issue as it indicates that maturational schedules themselves are shifting, and the effects of this change on adolescent mental health are currently unknown^120,121^. Our results show that incorporating maturational timing and maturity-related momentum into future studies, and ultimately, into clinical practice, could support targeted assessment and intervention: individuals on persistently low-hierarchy trajectories may require different forms and timing of support than those undergoing transient, high-momentum phases of change.

Overall, our study suggests two distinct forms of vulnerability. First, accelerated maturational timing is associated with persistent reduction in brain hierarchical organization. Second, periods of peak neurodevelopmental change define windows of heightened vulnerability, as seen in the deviation from the expected developmental trajectory in brain hierarchy in the decelerated group following the COVID-19 pandemic. Together, these findings indicate that accelerated and decelerated maturational timing do not simply represent symmetric deviations from a normative path, but reflect fundamentally distinct neurodevelopmental paths. Critically, these differences were only detectable by employing longitudinal trajectories modelling age-related patterns of neurodevelopment, highlighting the limitations of group-level, summarizing approaches for understanding maturation-dependent development.

These results highlight two converging threats to healthy adolescent brain development: the ongoing secular trend towards earlier puberty, which shifts entire maturational timelines, and global ecological crises that rapidly restructure living environments and social contexts. These changes are already linked to disturbances in mental health, cognition and overall well-being of adolescents across nations. Our results also emphasize the need to utilize objective maturity measurements in tailoring support for individuals according to their pubertal timelines, as evidently, pubertal timing is a critical dimension of adolescent brain vulnerability.

## Materials and Methods

### Participants

We employed a longitudinal study design with two waves of data collection. During the first wave (November 2017–July 2019), we assessed 87 adolescent female participants aged 11.60 to 15.97 (T1; mean age: 13.36, s.d.=0.97) before the COVID-19 pandemic. During the second wave (March 2023–November 2024), we reassessed participants in emerging adulthood (T2; 16.91–19.99 years; mean age: 18.24, s.d.=0.78; 64% retention rate) after COVID-19-related school closures were terminated in Hungary^64^. The mean within-participant interval between T1 and T2 data acquisition was 5.00 years (s.d.=0.54). T1 participants were recruited via school contacts and online advertisements from Budapest and Budaörs, Hungary. To assemble the T2 dataset, participants of the T1 sample were contacted through personal or school email, phone, or by using the contact information of a parent to repeat data acquisition procedures administered at T1.

All participants of the T1 sample underwent bone age screening. Participants were then classified into three non-overlapping maturity groups based on the difference between their CA and BA values, and approximately equal numbers were invited for laboratory testing from each group: decelerated (BA<CA-0.5, mean BA-CA: −1.01, s.d.=0.48; N=31), average (CA-0.5≤BA≤CA+0.5, mean BA-CA: −0.01, s.d.=0.28; N=28), and accelerated (BA>CA+0.5, mean BA-CA: 0.94, s.d.=0.36; N=28), following protocol established in earlier works^22,32^. Power calculations for a two-sample z-test at a power and effect size of 0.8 and a two-sided α of 0.05 yielded a required sample size of N ≥ 25 per group. To ensure a sufficient sample size, we aimed for N=30 per group. The sample size in the T1 maturity groups (N=28 and N=31) yields a power of at least 0.849, while the sample sizes in the T2 maturity groups (N=18 and N=19) yields a power of at least 0.67. As the chronological age range was not equal across maturity groups (decelerated: mean CA: 13.44, min.=12.10, max.=14.77, s.d.=0.87; average: mean CA: 13.26, min.=11.60, max.=15.97, s.d.=1.09; accelerated: mean CA: 13.37, min.=11.63, max.=15.07, s.d.=0.99), we chose not to divide the T1 dataset into age groups. We tracked menarche, a female-specific marker of pubertal development, as additional information via a self-report questionnaire. The decelerated maturity group exhibited the lowest percentage of postmenarche participants (48%; 75% in the average and accelerated groups) in the T1 sample, supporting the notion that ultrasonic bone age assessments provide a reliable measurement of maturity in adolescence.

Information on place of residence and parental education illustrates the high socioeconomic background of the participants. 84% of T1 participants lived in the capital city at the time of data acquisition, 60% attended an 8-year high school, and 31% a 6-year high school. Regarding parental education, 93% of mothers and 92% of fathers reported having a college or university degree. All information on parental education, place of residence, and percentage of post-menarche participants is reported in Tables S25, S26 and S27. A retrospective survey of T2 participants (64% response rate), assessing health and well-being during the pandemic, was conducted between May and October 2025. The survey included items on COVID-19 infection and vaccination history, the Post-COVID Symptom Scale for Children/Youth^122^ (PCSS-C/Y), as well as items from the Health Behaviour in School-aged Children (HBSC^123^; Hungarian translation^124^). A summary of responses is reported in Table S28.

We sorted T2 participants into the same maturity group as they were assigned to in T1, replicating the decelerated (N=19, mean CA: 18.53, min.=17.07, max.=19.85, s.d.=0.78), average (N=18, mean CA: 18.06, min.=16.91, max.=19.04, s.d.=0.62) and accelerated (N=19, mean CA: 18.12, min.=17.07, max.=19.99, s.d.=0.86) groups with balanced participant numbers. Repeated measurements included body height and weight measurements, and resting state EEG data collection. Bone age assessment was not repeated (see *Bone age assessment* in *Procedure* for further information).

To assess the possibility of selective retention, we compared participants who returned for T2 data acquisition with those with T1 data only, assessing differences in mean individual entropy production, chronological age, maturity (BA–CA), EEG recording length, number of removed EEG channels, age at menarche, parental education, place of residence, school type, and maturity group assignment. Comparisons were conducted on data collected at T1. The comparisons did not indicate selective retention. Results of these comparisons are reported in Table S29.

Written informed consent was obtained from all subjects and/or their parents prior to data collection in the T1 sample, and from all participants in the T2 sample. Participants were given gift cards (with an approximate value of EUR 15 each) for their attendance in both data acquisition periods. Parents and/or participants provided details about their demographic, educational, and medical histories. Only neurotypical participants were included in the current study. The Hungarian United Ethical Review Committee for Research in Psychology (EPKEB) approved the study (reference number 2017/114). All procedures were performed in compliance with the Declaration of Helsinki.

### Procedure

#### Bone age assessment

We conducted bone age estimations to test the maturity status of our participants at T1. Bone age assessments were conducted by trained assistants, overseen by a biological anthropologist, and took approximately 5-10 minutes per participant. Measurements were taken either at the schools of the participants or at the Research Centre for Sport Physiology at the Hungarian University of Sports Science, Budapest.

Bone age assessment procedures began with the measurement of full body and sitting height using a GPM Anthropometer (DKSH, Switzerland Ltd, Zurich, Switzerland) and body weight with a digital scale (Seca GmbH & Co. KG., Hamburg, Germany), followed by the estimation of bone age using the Sunlight BonAge System (Sunlight Medical Ltd, Tel Aviv, Israel), an ultrasonic device scanning the speed of sound between the distal radius and ulnar epiphysis, evaluating the degree of ossification within this structure, reflecting the level of maturation^125^. The BonAge System calculates bone age in years and months utilizing an algorithm that accounts for gender and ethnicity. Participants were instructed to place their left hand on the armrest of the scanner, between two transducers, where the ulnar side emits the ultrasound and the radial side receives the signals. Then, the device attached to the wrist, applying approximately 500g of pressure, and the ulnar transducer started to emit 750 kHz frequency ultrasound through the wrist for approximately 20 seconds. Measurements were repeated 5 times to ensure accuracy. Results of ultrasonic bone age assessment highly correlate with those of radiography^125,126^, however, the former does not expose the participants to ionizing radiation, making it a reliable and safe choice.

Bone age assessments are valid for 3 months; therefore, participants of the T1 sample were invited for the EEG data collection within this timeframe. Bone age assessment was repeated if a participant was unable to undergo laboratory testing within 3 months of the first screening.

As epiphyseal fusion at the distal radius and ulna is completed by 17–18 years in females^127^, this assessment was carried out only on the T1 sample. However, maturity group assignments were carried forward from T1 to T2 to examine whether pubertal timing would outline diverging developmental trajectories, and whether these differences would remain detectable in young adulthood.

#### Resting state EEG data acquisition

EEG data recording took place at the Laboratory for Psychological Research, Pázmány Péter Catholic University, Budapest, and followed established protocol^128,129^. EEG data acquisition was preceded by a brief guided relaxation and removal of all mobile electronic devices, such as phones or smartwatches, from the pockets and bodies of the participants. Two 300-s-long continuous EEG data segments were recorded using the 128-channel high-density EEG system HydroCel GSN 130 (Magstim EGI, Oregon, USA), a quick-application EEG sensor net, with the Net Station Acquisition software (version 5.4), at a 1000 Hz sampling rate, in a windowless, ventilated, and electromagnetically shielded room. We aimed to keep impedances below 50 KΩ across all channels. During the first 300-s-long recording segment, participants were seated and instructed to keep their eyes open, blink only minimally and remain still to reduce eye- and muscle movement-related artifacts^130^. We chose not to present a fixation point to the participants during the eyes-open recording segment to avoid directing their visual attention and therefore introducing fixation-related potentials into our data^131^. An identical protocol was followed during the second recording segment, except participants were instructed to keep their eyes closed during the entire 300-s-long session. The latter instruction was explained to participants during a short break between the first and second 300-s-long data recording session, while participants were still seated and the sensor net was not removed. The beginning and end of the eyes-open and eyes-closed sequences were manually annotated in the Net Station Acquisition interface. The recording environment and procedure was uniform in all cases in both T1 and T2 samples, minimizing the confounding effects of external stimuli.

Spectral analysis has previously been performed on a set of EEG recordings^32^ with considerable overlap with the T1 sample of the present study (N=56, aged 12-13 years), verifying that the EEG recordings demonstrate the well-known and expected difference of alpha power between the eyes-closed and eyes-open recordings, and also the established age-related changes of the spectra^132,133^, providing evidence for the reproducibility and validity of our EEG measurements. In this previous work, we have also established that the eyes-closed recordings are the best candidates on which to carry out entropy production calculations, as this resting state condition is dominated by the alpha frequency component of the EEG spectrum in both adolescents^132^ and adults^134^, reflecting spontaneous brain activity more accurately than the eyes-open condition^135^. Furthermore, we were unable to demonstrate significant maturity-related modifications on the eyes-open dataset in terms of entropy production. Based on this information, we focused the current analysis on the eyes-closed recordings and the alpha frequency range.

The EEG data collection period of T1 spanned from November 2017 to July 2019, and that of T2 from March 2023 to November 2024. Resting state EEG data acquisition procedure was identical at both data acquisition waves.

#### EEG data preprocessing

EEG preprocessing was performed in EEGLAB^136^ (version: 2021.1) in a MATLAB environment (The MathWorks Inc., version: R2021a, academic license). First, a high-pass filter at 1 Hz, with a transition bandwidth of 1 Hz, a passband edge of 1 Hz, and a cutoff frequency of −6 dB 0.5 Hz, was applied to filter out low-frequency noise and direct-current offset using the default finite impulse response (FIR) filter (version: 2.4) within EEGLAB. Then, a 40 Hz low-pass filter with a transition bandwidth of 10 Hz, a passband edge of 40 Hz, and a cutoff frequency of −6 dB 45 Hz was applied via the same filtering algorithm. Following filtering, channel locations were imported into EEGLAB using the layout map provided by Magstim EGI. Noisy channels were selected by an experienced rater after visual inspection, then removed from the recordings manually. The number of removed channels per EEG recording ranged from 0 to 15 in the T1 sample and from 0 to 8 in the T2 sample. The distribution of removed channels within each maturity group is reported in Table S30. Automatic epoch removal was performed with the Clean Rawdata plugin (version: 2.4), which rejects 1-s long sliding windows (66% overlap) whose variance exceeds 20 times the standard deviation of the calibration data. After this step, the average length of recordings was 282 seconds in the T1 sample (s.d.=27.59) and 285 seconds in the T2 sample (s.d.=24.14). Finally, recordings were re-referenced to average reference. Preprocessing steps were uniform for all recordings.

#### Leading Eigenvector Dynamics Analysis

Following preprocessing steps, we applied the Leading Eigenvector Dynamics Analysis^62^ (LEiDA) algorithm to each preprocessed EEG recording. LEiDA was adapted to EEG data by our group in order to capture the instantaneous global phase relationships of EEG signals over the cortex, ranging from in-phase to counter-phase, represented as topographically distinct activity patterns referred to here as brain states. The algorithm steps were followed exactly as previously described^32^ and were conducted in MATLAB (version: R2021a, academic license).

Preprocessed EEG recordings were first bandpass filtered to the alpha range (8-12 Hz) using a 6th-order Butterworth filter with zero phase shift. The data were then downsampled to 100 Hz (10 ms intervals). Signals were subsequently detrended and demeaned, and the instantaneous phase for all channels at each time point *t* was extracted via Hilbert-transformation.

In order to obtain 128×128 coherence matrices at each time point *t* and for every subject, compensating for the noisy channels removed during preprocessing, we followed a two-step data patching process. First, a maturity group-specific list of noisy channels was compiled separately for the T1 and T2 samples. For the *n* remaining “good channels”, common to all participants in a group, instantaneous phase coherence *PC_xy,t_* for each pair of electrodes was calculated at each time point *t* using the following formula:

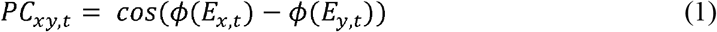

where *ϕE_x,t_* and *ϕE_y,t_* stands for the instantaneous phase of electrodes *x* and *y* at time point *t*. As a result, a sequence of *n*x*n* phase coherence matrices was computed at 10-millisecond intervals for each subject. At each time point *t*, the leading eigenvector (1x*n*) of each coherence matrix was computed, summarizing the instantaneous phase coherence of all electrodes. We applied the built-in *k*-means clustering algorithm in MATLAB to all leading eigenvectors, enabling us to assign a cluster label to all sampled timepoints of each EEG recording. After this first clustering process, phase values missing due to channel rejection were reconstructed in the last stage of patching. For each bad channel *x* at each time point *t* assigned to a particular cluster, missing phase values were reconstructed by randomly drawing from the pool of phase values measured at electrode *x* in all other EEG recordings within the same maturity group, restricted to time points labeled with the same cluster.

Repeating this process across all removed channels in each EEG recording and each maturity group in both the T1 and T2 samples yielded a collection of complete phase vectors (1×128) and phase coherence matrices (128×128). A new set of leading eigenvectors (1×128) were then derived from the completed matrices, matching the original phase vectors in size but denoting relative, not absolute, phase. Patching steps were conducted separately for the T1 and T2 samples.

Following patching, the polarity-invariant patterns of phase coherence common across EEG recordings, referred to as brain states, were identified based on the cosine similarity of leading eigenvectors for the T1 and T2 samples separately. First, leading eigenvectors were pooled from all sampled timepoints from all EEG recordings, and clustered using a modified *k*-means algorithm in which each vector is assigned to a cluster, representing a brain state, labeled accordingly. Sample-specific brain states are represented in Fig. 6. The application of LEiDA to resting state EEG recordings allowed us to parse each EEG recording into a sequence of brain states and to quantify the temporal asymmetry of switching within each brain state pair, revealing the level of hierarchical organization in EEG recordings.

**Fig. 6:**
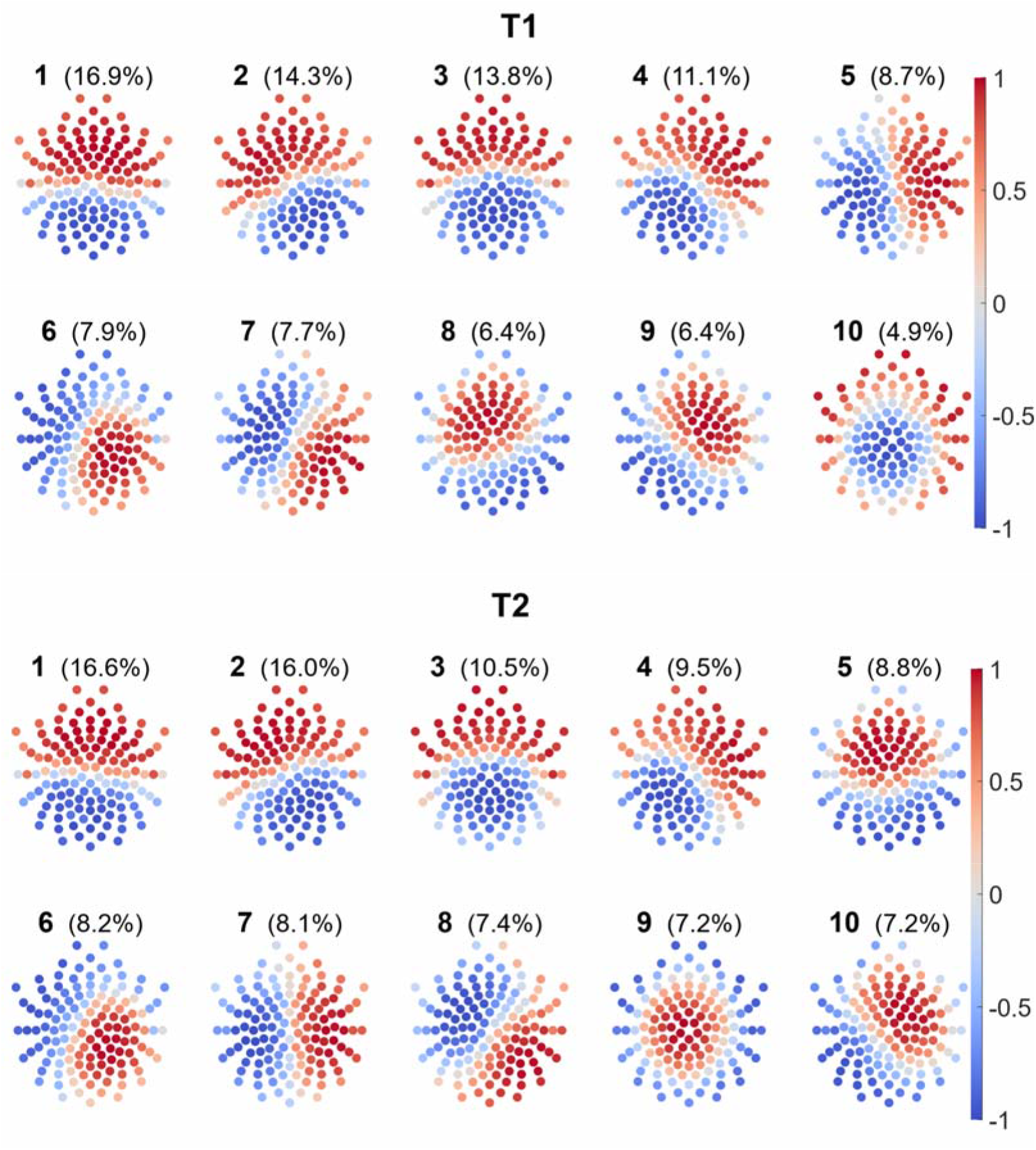
Brain states of the T1 and T2 samples. The figure presents two sets of 10 brain states for the T1 and T2 samples, as indicated above each set. Brain states are derived from eyes-closed resting state EEG recordings by applying LEiDA^62^ to clean data in the alpha frequency band (8-12 Hz). For each recording, the instantaneous phase of each channel is extracted via Hilbert transformation, then instantaneous phase coherence matrices and the corresponding leading eigenvectors are computed at each time point *t* (10-millisecond intervals). Within each sample, leading eigenvectors from each time point *t* are pooled, then clustered via a *k*-means algorithm, assigning each vector to one of *k* clusters, each representing a brain state. For each sample, brain states are arranged in two rows and ranked from 1 to 10 in descending order according to the fraction of recording time spent in each state, reflected by percentages accompanying rank labels. Both color scales to the right of the sample-specific brain state sets depict values on a −1 to +1 scale consistent with cosine similarity, from +1 (red; in phase) to −1 (blue; counter-phase), denoting the spectrum of possible phase alignments. By applying LEiDA, we defined each EEG recording as a sequence of brain states, enabling the quantification of temporal asymmetries between state pair transitions indicating the level of hierarchical organization.

A sensitivity analysis using a common clustering solution across waves indicated that merging T1 and T2 samples introduces cross-wave noise that attenuates otherwise detectable effects, supporting the use of wave-specific clustering. Results of this analysis are reported on the OSF repository of this project, accessible at https://osf.io/9qtv4/overview?view_only=a2d0eed5ea644c518cf34f36ef824ca4.

In the present study, we chose to conduct analyses using 10 brain states to provide sufficient resolution of the data while also maintaining computational efficiency, considering the large number of data points per sample. Previously, we have shown that analysis results are consistent across 6, 8, 10 and 12 brain states^32^.

### Mean individual entropy production calculation

To estimate the level of hierarchical organization in EEG recordings, the temporal asymmetry of switching within each brain state pair was quantified through the calculation of entropy production. First, a transition count table *N*_ij_ was constructed based on each subject-specific brain state sequence, including the number of forward (_i,j_) and reverse (_j,i_) transitions between each brain state pair. The diagonal entries of tables representing self-transitions, or remaining in the same state, were removed. To prevent zero-probability transitions in entropy production calculations, we applied Laplace smoothing^137^ to all transition counts (*N”*_ij_=*N*_ij_+1). (The effect of this smoothing on entropy production calculation was later neutralized via a noise floor correction procedure.) Then, transition count tables were transformed into transition probability tables for each subject, where transition probabilities (*P*) were calculated as *P*_ij_ = *N”*_ij_/Σ_ij_*N”*_ij_. Entropy production was then calculated based on subject-specific state transition probability tables via the Kullback-Leibler divergence:

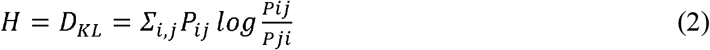

where *H* denotes entropy production calculated as the Kullback-Leibler Divergence (*D_KL_*), *P*_ij_ the probability of transitioning from brain state *i* to *j*, and *P*_ji_ the probability of reverse transitions (from brain state *j* to *i*).

To calculate mean entropy production values and accompanying variance for each participant, 1000 synthetic transition trajectories were generated based on the observed subject-specific transition probability tables, matching their length, utilizing two functions within the Economics Toolbox (version: 9.10.0.1684407) in MATLAB: ‘dtmc’ to generate discrete-time Markov chains and ‘simulate’ to simulate Markov chain state walks. Then, we applied the Kullback-Leibler divergence as specified earlier to the synthetic trajectories, resulting in 1000 observed entropy production data points for each subject, from which an individual mean entropy production value and its accompanying variance were calculated.

To offset the effects of Laplace smoothing and that of inflation introduced by the finite length of subject-specific trajectories to entropy production calculations, a modified version of the noise floor correction described by Lynn and colleagues^53^ was performed. To this end, a symmetric (null) transition table was first obtained for each subject by averaging the original observed transition probability table with its transpose, then 1000 synthetic length-matched null trajectories were generated and 1000 null entropy production data points calculated through the same steps as described above. This procedure revealed nonzero entropy production values, consistent finite data length- and smoothing-induced inflation. Noise floor-corrected entropy production values were obtained by calculating the mean of the 1000 null entropy production data points, then subtracting the null mean from the 1000 observed entropy production data points per subject. The procedure described here accounted for the temporal characteristics of subject-specific brain dynamics. Finally, the mean and variance of the 1000 noise-floor corrected entropy production values per subject were calculated using the following formula:

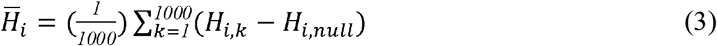

where *H̄_i_* is the mean entropy production of subject *i*, *H_i,k_*, is the entropy production for the *k*^th^ synthetic trajectory, and *H_i,null_*, is the mean null entropy production for subject *i*. The mean entropy production values were then utilized in the Gompertz function fitting procedure as described below. All calculations were performed in MATLAB (version: R2021a, academic license).

### Gompertz function fitting

Variations of the Gompertz function have been used to characterize multiple facets of brain development, including longitudinal white matter development^44^, regional myelin maturation and cortical development during infancy^45,46^, normative brain volume increase^47^, and maturation of the adolescent sleep EEG^48,49^. The three-parameter version – upper asymptote (α), shift parameter (β), and growth rate (γ) – captures key features of human brain development: asymptotic growth toward a developmental ceiling^47^, asymmetric dynamics with rapid early change followed by gradual tapering^34^, and a quantifiable rate of change^138^. In terms of neurodevelopmental studies specifically utilizing EEG recordings, Feinberg, Campbell and colleagues routinely used the Gompertz function to describe indices of cortical maturation derived from adolescent EEG, demonstrating the biological interpretability of the shape and parameters of the function^48,49,139,140^ in the context of adolescent cortical development, supporting its applicability in the current framework of hierarchical brain organization and its developmental dynamics. A sensitivity analysis comparing bootstrapped Gompertz and linear fits at T1 via Kolmogorov-Smirnov tests confirmed that the theoretically and practically grounded choice of the Gompertz function was not made at a cost in goodness of fit (Fig. S2, Table S31). Gompertz function fitting and subsequent statistical analyses were performed in MATLAB (version: R2021a, academic license), using a custom pipeline developed by our group.

First, a random number generator was initialized with the fixed seed (1) to ensure reproducibility of results. Then, the anonymized identification (ID) code, maturity group assignment, chronological age values, and mean individual entropy production values were imported into MATLAB. The entire CA range on which to perform the Gompertz function fitting was defined as a column vector of CA grid values from the observed global minimum (11.60) to global maximum (19.99) in 0.01 year increments.

Following data import and initial settings, a bootstrap loop (n=2000) was set up to perform the Gompertz function fitting on bootstrapped data points within T1 and T2 separately, allowing the estimation of the distribution of entropy production at each CA grid point.

In each loop, CA and entropy production value pairs of the starting dataset were resampled with replacement to match the sample size of the starting dataset, utilizing the built-in ‘randi’ function in MATLAB. After each bootstrap replication, the Gompertz function fitting procedure was performed on the resampled data. We defined the Gompertz function as

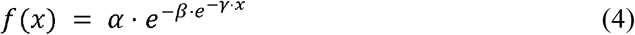

where *α* is the asymptote or maximum value of the function, *β* is the displacement parameter defining the horizontal shift of the function, *γ* is the growth rate, setting the scaling along the y-axis, and *e* is Euler’s number. Lower and upper parameter bounds for *α, β* and *γ* were set as (0.008, 0.001, 0.01) and (0.3, 100, 4), respectively. α parameter bounds were set to allow the curve sufficient flexibility to capture potential growth patterns dictated by the data. β and γ bounds were determined via iterative parameter space exploration of the complete datasets (both T1 and T2 samples, and maturity groups pre-bootstrap), selecting the parameter set that minimized the sum of squared residuals (SSR) across all subgroups. These bounds were then held constant during bootstrap resampling. All fitted parameters remained within the specified bounds in both initial explorative fits and bootstrap resamples. Optimization was performed using the ‘lsqcurvefit’ function, a nonlinear least-squares solver in the Optimization Toolbox within MATLAB, with a function tolerance of 10^-6^, combined with a MultiStart function of the Global Optimization Toolbox, which determines the number of random starting points within the ‘lsqcurvefit’ function. MultiStart was employed in the Gompertz function fitting process to decrease the likelihood of convergence on local minima, and to ensure a more robust search for the best possible fit on the data by generating 500 initial guesses within the lower and upper bounds, retaining the parameter combination which yielded the lowest SSR. Initial parameters were chosen randomly within the parameter bounds at each fitting iteration. The best parameter combination, defined as those yielding the lowest SSR, was saved in each bootstrap iteration. All fitting iterations converged successfully.

All described steps were performed separately on the decelerated, average and accelerated maturity group-specific dataset, with T1 and T2 data points handled separately, and also the complete T1 and T2 datasets, in which maturity groups were merged, resulting in a total of 8 sub-datasets. Bootstrapping and subsequent Gompertz function fitting was performed 2000 times, resulting in 2000 parameter combinations and 2000 function values in each sub-dataset over the global CA span. 2000 bootstrap replications are recommended in previous literature^141,142^ to sufficiently ensure that error introduced through randomization remains small relative to the underlying sampling error^143^, yielding stable estimates of uncertainty. To aid visualization, a mean Gompertz function and an 83.5% confidence interval were calculated within each sub-dataset by averaging over the 2000 function values, then finding their 8.25^th^ and 91.75^th^ percentiles at each CA point. We chose to depict 83.5% confidence intervals on our figures as their non-overlap effectively signifies statistically meaningful differences at the level of <0.05^144^.

### Statistical comparison of Gompertz functions

Pairwise comparisons of bootstrapped Gompertz functions were conducted over predefined CA subintervals utilizing an area-under-the-curve (AUC) approach. Similar integrated area approaches have been previously used to measure non-overlap of a sample and a reference distribution^145^, to compare survival curves^146^ and assess survival benefit^147^, to analyze urine estradiol and testosterone levels in adolescents^10^, and to test the capacity of a quantitative marker for treatment selection^148^.

For comparisons conducted within the T1 sample (average vs. accelerated; average vs. decelerated; decelerated vs. accelerated), the comparison domain was 11.60–15.97 years, partitioned into predefined, 0.25-year-wide CA windows between 11.60 and 15.75 years, and a final 0.21 year-wide window from 15.76 to 15.97 years, with a 0.01 year gap between each successive window to avoid pseudoreplication. Contrasts of T1-based projections and T2 bootstrapped Gompertz functions (all maturity groups pooled; accelerated; average; decelerated) were conducted over the range of 16.91-19.99 years, spanning the CA range of the T2 sample, partitioned into 0.25-year-wide CA windows between 16.91 and 19.76 years, and a 0.22-year-wide CA window between 19.77 and 19.99 years. First, within each predefined CA window, we computed the pointwise difference between the two bootstrapped Gompertz functions as

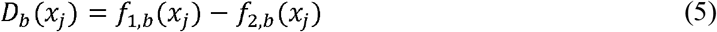

where *D_b_(x_j_)* is the pointwise difference of functions at CA point *x_j_*, *f_1,b_* and *f_2,b_* are the bootstrapped function values, and *b* is the bootstrap replicate index. This step produced 2000 difference functions of the same length as the predefined CA windows. Then, we computed the area under each difference function by numerical (trapezoidal) integration utilizing the built-in ‘trapz’ function in MATLAB, yielding 2000 AUC values per predefined CA window. From these 2000 AUC values, we calculated the window specific mean and 95% confidence interval defined as the 2.5th and 97.5th percentile of the window-specific AUC distribution. A window was considered to show significant divergence if the two-sided 95% confidence interval of the mean area did not include 0.

### Slope calculation

To compare differences in the rate of change within the T1 sample (average vs. accelerated; average vs. decelerated; decelerated vs. accelerated), slopes were calculated over 11.60–15.97 years, partitioned into 0.25-year-wide CA windows between 11.60 and 15.75 years, and a 0.21 year-wide window from 15.76 to 15.97 years. Instantaneous slopes were obtained from the numerical derivative of each bootstrap-derived Gompertz function, computed using the built-in MATLAB function ‘gradient’, and then averaged across the specified CA windows.

To statistically compare slopes across maturity groups, we computed the difference of window-specific slopes between two groups as

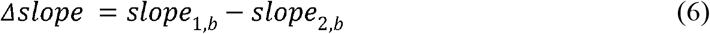

Following this, we calculated the mean Δslope and the two-sided 95% CI of each CA window and flagged a statistically significant difference between slopes if the CI did not include 0.

### Sensitivity analyses

To assess the robustness of our results to window width, we repeated all trajectory and slope comparisons using 0.50-year, 1.0-year, and 1.5-year non-overlapping CA windows. CA intervals for each windowing approach are detailed in Table 1. Sensitivity analyses supported the robustness of the results reported in the main text. Sensitivity analysis results are presented in the Supplementary Material (Tables S15–S24).

**Table 1.** Windowing approaches and CA intervals of sensitivity analyses assessing the robustness of trajectory and slope comparisons across adolescence, and of projected vs. measured trajectories over the T2 age range. Window widths are noted in parentheses where the final window differs from the nominal width.

| **Windowing approach** | **CA intervals T1** | **CA intervals T2** |
|---|---|---|
| 0.50 | 11.60–12.10 | 16.91–17.41 |
| 0.50 | 12.11–12.61 | 17.42–17.92 |
| 0.50 | 12.62–13.12 | 17.93–18.43 |
| 0.50 | 13.13–13.63 | 18.44–18.94 |
| 0.50 | 13.64–14.14 | 18.95–19.45 |
| 0.50 | 14.15–14.65 | 19.46–19.99 (0.53 years) |
| 0.50 | 14.66–15.16 |  |
| 0.50 | 15.17–15.67 |  |
| 0.50 | 15.68–15.97 (0.29 years) |  |
| 1.0 | 11.60–12.60 | 16.91–17.91 |
| 1.0 | 12.61–13.61 | 17.92–18.92 |
| 1.0 | 13.62–14.62 | 18.93–19.99 (1.06 years) |
| 1.0 | 14.63–15.97 (1.34 years) |  |
| 1.5 | 11.60–13.10 | 16.91–18.41 |
| 1.5 | 13.11–14.61 | 18.42–19.99 (1.57 years) |
| 1.5 | 14.62–15.97 (1.35 years) |  |

### Inverse-variance weighted average calculation

To mitigate the biasing effect of heterogeneity and high subject-specific variance on the mean, average entropy production and variance of maturity groups were calculated via inverse-variance weighting as

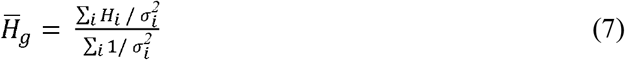

where *H̄_g_* denotes the inverse-variance weighted average of the specified maturity group, *H_i_* the mean individual entropy production and 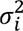 the observed variance of subject *i*. The inverse-variance weighted group variance was calculated as

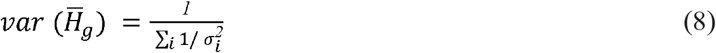

The upper and lower values of 95% confidence intervals accompanying inverse-variance weighted averages (as shown in Fig. 4) were calculated as 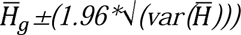. Calculations of inverse-variance weighted averages and group variances were carried out in MATLAB (version: R2021a, academic license).

### Two-sample z-tests

Statistical comparisons on the group level were performed via two-sample z-tests^149^ using the following formula:

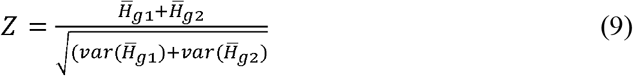

The level of significance was determined as

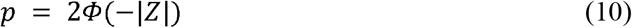

Where *ϕ* denotes the standard normal cumulative distribution function.

Significant results obtained from the two-sample z-tests were reported with a Hedges’ *g* effect size estimations. Two-sample z-tests and effect size estimations were carried out in Microsoft Excel (version: 2507).

## Data availability

All data needed to evaluate the conclusions in the paper are present in the paper and the accompanying Supplementary Material. Individual-level data are accessible at the OSF repository of this project (https://osf.io/n2xz5?view_only=a2d0eed5ea644c518cf34f36ef824ca4).

## Code availability

The custom MATLAB (version R2021a) code used for entropy production calculations, Gompertz function fitting and bootstrap analyses is available from the corresponding author upon reasonable request.

## Acknowledgements

The authors thank K. Utczás and Zs. Tróznai for bone age data collection and analysis, Gy. Oláh and J. V. Dénes for organising the participant cohort, and L. J. Fehér for EEG data collection.

The project was supported by the National Research, Development and Innovation Office of Hungary (Grant K-134370 to I.K.) and by the Hungarian Research Network (HUN-REN-ELTE-PPKE Adolescent Development Research Group). It was made in the framework of the PPKE-BTK-KUT-23-1 project, with the support and funding provided by the Faculty of Humanities and Social Sciences of Pázmány Péter Catholic University. The funders had no role in study design, data collection and analysis, decision to publish or preparation of the manuscript.

G.D. is supported by Grant PID2022-136216NB-I00 funded by MICIU/AEI/10.13039/ 501100011033 and by “ERDF A way of making Europe”, ERDF, EU, Project NEurological MEchanismS of Injury, and Sleep-like cellular dynamics (NEMESIS) (ref. 101071900) funded by the EU ERC Synergy Horizon Europe, and AGAUR research support grant (ref. 2021 SGR 00917) funded by the Department of Research and Universities of the Generalitat of Catalunya.

M.L.K. is supported by the Centre for Eudaimonia and Human Flourishing (funded by the Pettit, Carlsberg and Cillo Foundations) and Center for Music in the Brain (funded by the Danish National Research Foundation, DNRF117, The Lundbeck Foundation R469-2024-1573 and Købmand Herman Sallings Fond).

## Author contributions

H.S. collected the data, conducted formal analyses, created figures and wrote the first draft. I.K., F.G., and H.S. conceived the methodology and interpreted the data. M.L.K. and G.D. contributed theoretical and methodological framework. I.K. designed the paradigm, secured funding and supervised the project. All authors discussed the results, revised the manuscript, and approved the final version for submission.

