## Supplemental Data 1 for "Pubertal timing is a critical dimension of adolescent brain vulnerability"

Ilona Kovács, HUN-REN-ELTE-PPCU Adolescent Development Research Group, Budapest, 1088 Hungary

### Accelerated maturity is associated with persistent reduction in hierarchical brain organization - Tables S1-S6

| **CA interval** | **Mean AUC** | **95% CI (lower)** | **95% CI (upper)** | **Significant (1: yes; 0: no)** |
| --- | --- | --- | --- | --- |
| 11.60–11.85 | -0.00113 | -0.00409 | 0.00168 | 0 |
| 11.86–12.11 | -0.00130 | -0.00393 | 0.00123 | 0 |
| 12.12–12.37 | -0.00147 | -0.00382 | 0.00078 | 0 |
| 12.38–12.63 | -0.00160 | -0.00356 | 0.00030 | 0 |
| 12.64–12.89 | -0.00185 | -0.00375 | -0.00012 | 1 |
| 12.90–13.15 | -0.00206 | -0.00380 | -0.00047 | 1 |
| 13.16–13.41 | -0.00228 | -0.00404 | -0.00064 | 1 |
| 13.42–13.67 | -0.00250 | -0.00437 | -0.00076 | 1 |
| 13.68–13.93 | -0.00274 | -0.00487 | -0.00082 | 1 |
| 13.94–14.19 | -0.00298 | -0.00545 | -0.00080 | 1 |
| 14.20–14.45 | -0.00324 | -0.00614 | -0.00077 | 1 |
| 14.46–14.71 | -0.00337 | -0.00659 | -0.00072 | 1 |
| 14.72–14.97 | -0.00363 | -0.00742 | -0.00066 | 1 |
| 14.98–15.23 | -0.00406 | -0.00863 | -0.00062 | 1 |
| 15.24–15.49 | -0.00435 | -0.00963 | -0.00053 | 1 |
| 15.50–15.75 | -0.00464 | -0.01058 | -0.00043 | 1 |
| 15.76–15.97 | -0.00394 | -0.00921 | -0.00026 | 1 |

**Table S1.** Chronological age (CA) intervals, mean area-under-the-curve (AUC) values, and 95% confidence interval (CI) bounds for the comparison of the average vs. accelerated maturity groups at T1. Significance of each CA window at a level of α<0.05 (two-sided) is indicated in the last column.

| **CA interval** | **Mean AUC** | **95% CI (lower)** | **95% CI (upper)** | **Significant (1: yes; 0: no)** |
| --- | --- | --- | --- | --- |
| 11.60–11.85 | 0.00299 | 0.00022 | 0.00563 | 1 |
| 11.86–12.11 | 0.00287 | 0.00034 | 0.00538 | 1 |
| 12.12–12.37 | 0.00274 | 0.00039 | 0.00506 | 1 |
| 12.38–12.63 | 0.00249 | 0.00046 | 0.00450 | 1 |
| 12.64–12.89 | 0.00244 | 0.00048 | 0.00442 | 1 |
| 12.90–13.15 | 0.00227 | 0.00047 | 0.00420 | 1 |
| 13.16–13.41 | 0.00209 | 0.00026 | 0.00403 | 1 |
| 13.42–13.67 | 0.00190 | -0.00008 | 0.00406 | 0 |
| 13.68–13.93 | 0.00170 | -0.00061 | 0.00411 | 0 |
| 13.94–14.19 | 0.00149 | -0.00119 | 0.00426 | 0 |
| 14.20–14.45 | 0.00128 | -0.00194 | 0.00465 | 0 |
| 14.46–14.71 | 0.00101 | -0.00261 | 0.00483 | 0 |
| 14.72–14.97 | 0.00080 | -0.00338 | 0.00531 | 0 |
| 14.98–15.23 | 0.00061 | -0.00444 | 0.00615 | 0 |
| 15.24–15.49 | 0.00038 | -0.00541 | 0.00674 | 0 |
| 15.50–15.75 | 0.00014 | -0.00643 | 0.00741 | 0 |
| 15.76–15.97 | -0.00005 | -0.00599 | 0.00640 | 0 |

**Table S2.** CA intervals, mean AUC values, and 95% CI bounds for the comparison of the average vs. decelerated maturity groups at T1. Significance of each CA window at a level of α<0.05 (two-sided) is indicated in the last column.

| **CA interval** | **Mean AUC** | **95% CI (lower)** | **95% CI (upper)** | **Significant (1: yes; 0: no)** |
| --- | --- | --- | --- | --- |
| 11.60–11.85 | 0.00186 | 0.00007 | 0.00344 | 1 |
| 11.86–12.11 | 0.00158 | -0.00008 | 0.00311 | 0 |
| 12.12–12.37 | 0.00127 | -0.00024 | 0.00274 | 0 |
| 12.38–12.63 | 0.00090 | -0.00046 | 0.00223 | 0 |
| 12.64–12.89 | 0.00059 | -0.00076 | 0.00188 | 0 |
| 12.90–13.15 | 0.00021 | -0.00115 | 0.00146 | 0 |
| 13.16–13.41 | -0.00019 | -0.00159 | 0.00107 | 0 |
| 13.42–13.67 | -0.00060 | -0.00206 | 0.00075 | 0 |
| 13.68–13.93 | -0.00104 | -0.00266 | 0.00045 | 0 |
| 13.94–14.19 | -0.00149 | -0.00334 | 0.00029 | 0 |
| 14.20–14.45 | -0.00196 | -0.00415 | 0.00013 | 0 |
| 14.46–14.71 | -0.00235 | -0.00479 | -0.00001 | 1 |
| 14.72–14.97 | -0.00283 | -0.00564 | -0.00011 | 1 |
| 14.98–15.23 | -0.00345 | -0.00683 | -0.00025 | 1 |
| 15.24–15.49 | -0.00397 | -0.00796 | -0.00034 | 1 |
| 15.50–15.75 | -0.00450 | -0.00905 | -0.00035 | 1 |
| 15.76–15.97 | -0.00399 | -0.00811 | -0.00028 | 1 |

**Table S3**. CA intervals, mean AUC values, and 95% CI bounds for the comparison of the accelerated vs. decelerated maturity groups at T1. Significance of each CA window at a level of α<0.05 (two-sided) is indicated in the last column.

| **CA interval** | **Δslope** | **95% CI (lower)** | **95% CI (upper)** | **Significant (1: yes; 0: no)** |
| --- | --- | --- | --- | --- |
| 11.60–11.85 | -0.00250 | -0.00720 | 0.00362 | 0 |
| 11.86–12.11 | -0.00264 | -0.00782 | 0.00347 | 0 |
| 12.12–12.37 | -0.00279 | -0.00834 | 0.00333 | 0 |
| 12.38–12.63 | -0.00294 | -0.00897 | 0.00325 | 0 |
| 12.64–12.89 | -0.00309 | -0.00972 | 0.00311 | 0 |
| 12.90–13.15 | -0.00324 | -0.01031 | 0.00301 | 0 |
| 13.16–13.41 | -0.00340 | -0.01106 | 0.00290 | 0 |
| 13.42–13.67 | -0.00355 | -0.01171 | 0.00279 | 0 |
| 13.68–13.93 | -0.00370 | -0.01237 | 0.00265 | 0 |
| 13.94–14.19 | -0.00385 | -0.01294 | 0.00260 | 0 |
| 14.20–14.45 | -0.00399 | -0.01357 | 0.00255 | 0 |
| 14.46–14.71 | -0.00413 | -0.01409 | 0.00248 | 0 |
| 14.72–14.97 | -0.00427 | -0.01459 | 0.00240 | 0 |
| 14.98–15.23 | -0.00440 | -0.01508 | 0.00235 | 0 |
| 15.24–15.49 | -0.00453 | -0.01558 | 0.00230 | 0 |
| 15.50–15.75 | -0.00464 | -0.01600 | 0.00227 | 0 |
| 15.76–15.97 | -0.00475 | -0.01636 | 0.00226 | 0 |

**Table S4**. CA intervals, mean $\Delta$slope values, and 95% CI bounds for the comparison of the average vs. accelerated maturity groups at T1. Significance of each CA window at a level of α<0.05 (two-sided) is indicated in the last column.

| **CA interval** | **Δslope** | **95% CI (lower)** | **95% CI (upper)** | **Significant (1: yes; 0: no)** |
| --- | --- | --- | --- | --- |
| 11.60–11.85 | -0.00162 | -0.00651 | 0.00320 | 0 |
| 11.86–12.11 | -0.00187 | -0.00703 | 0.00349 | 0 |
| 12.12–12.37 | -0.00211 | -0.00759 | 0.00388 | 0 |
| 12.38–12.63 | -0.00233 | -0.00828 | 0.00427 | 0 |
| 12.64–12.89 | -0.00253 | -0.00894 | 0.00461 | 0 |
| 12.90–13.15 | -0.00271 | -0.00977 | 0.00508 | 0 |
| 13.16–13.41 | -0.00287 | -0.01051 | 0.00551 | 0 |
| 13.42–13.67 | -0.00301 | -0.01133 | 0.00620 | 0 |
| 13.68–13.93 | -0.00314 | -0.01206 | 0.00674 | 0 |
| 13.94–14.19 | -0.00324 | -0.01296 | 0.00729 | 0 |
| 14.20–14.45 | -0.00334 | -0.01374 | 0.00771 | 0 |
| 14.46–14.71 | -0.00341 | -0.01462 | 0.00820 | 0 |
| 14.72–14.97 | -0.00347 | -0.01547 | 0.00885 | 0 |
| 14.98–15.23 | -0.00352 | -0.01621 | 0.00939 | 0 |
| 15.24–15.49 | -0.00356 | -0.01684 | 0.00988 | 0 |
| 15.50–15.75 | -0.00358 | -0.01750 | 0.01037 | 0 |
| 15.76–15.97 | -0.00359 | -0.01812 | 0.01091 | 0 |

**Table S5**. CA intervals, mean $\Delta$slope values, and 95% CI bounds for the comparison of the average vs. decelerated maturity groups at T1. Significance of each CA window at a level of α<0.05 (two-sided) is indicated in the last column.

| **CA interval** | **Δslope** | **95% CI (lower)** | **95% CI (upper)** | **Significant (1: yes; 0: no)** |
| --- | --- | --- | --- | --- |
| 11.60–11.85 | -0.00411 | -0.00723 | -0.00011 | 1 |
| 11.86–12.11 | -0.00451 | -0.00781 | -0.00040 | 1 |
| 12.12–12.37 | -0.00489 | -0.00837 | -0.00060 | 1 |
| 12.38–12.63 | -0.00527 | -0.00886 | -0.00086 | 1 |
| 12.64–12.89 | -0.00562 | -0.00950 | -0.00095 | 1 |
| 12.90–13.15 | -0.00595 | -0.01023 | -0.00110 | 1 |
| 13.16–13.41 | -0.00627 | -0.01117 | -0.00113 | 1 |
| 13.42–13.67 | -0.00656 | -0.01202 | -0.00117 | 1 |
| 13.68–13.93 | -0.00684 | -0.01293 | -0.00114 | 1 |
| 13.94–14.19 | -0.00709 | -0.01386 | -0.00107 | 1 |
| 14.20–14.45 | -0.00733 | -0.01482 | -0.00106 | 1 |
| 14.46–14.71 | -0.00755 | -0.01584 | -0.00094 | 1 |
| 14.72–14.97 | -0.00774 | -0.01677 | -0.00078 | 1 |
| 14.98–15.23 | -0.00792 | -0.01758 | -0.00064 | 1 |
| 15.24–15.49 | -0.00808 | -0.01832 | -0.00051 | 1 |
| 15.50–15.75 | -0.00822 | -0.01912 | -0.00037 | 1 |
| 15.76–15.97 | -0.00834 | -0.01973 | -0.00027 | 1 |

**Table S6**. CA intervals, mean $\Delta$slope values, and 95% CI bounds for the comparison of the accelerated vs. decelerated maturity groups at T1. Significance of each CA window at a level of α<0.05 (two-sided) is indicated in the last column.

### Deviation in hierarchical brain organisation following the pandemic - Table S7

| **CA interval** | **Mean AUC** | **95% CI (lower)** | **95% CI (upper)** | **Significant (1: yes; 0: no)** |
| --- | --- | --- | --- | --- |
| 16.91–17.16 | 0.00463 | 0.00050 | 0.01050 | 1 |
| 17.17–17.42 | 0.00507 | 0.00050 | 0.01169 | 1 |
| 17.43–17.68 | 0.00509 | 0.00042 | 0.01197 | 1 |
| 17.69–17.94 | 0.00555 | 0.00038 | 0.01325 | 1 |
| 17.95–18.20 | 0.00556 | 0.00035 | 0.01356 | 1 |
| 18.21–18.46 | 0.00605 | 0.00025 | 0.01500 | 1 |
| 18.47–18.72 | 0.00605 | 0.00016 | 0.01525 | 1 |
| 18.73–18.98 | 0.00657 | 0.00008 | 0.01683 | 1 |
| 18.99–19.24 | 0.00655 | -0.00003 | 0.01684 | 0 |
| 19.25–19.50 | 0.00681 | -0.00011 | 0.01754 | 0 |
| 19.51–19.76 | 0.00737 | -0.00027 | 0.01918 | 0 |
| 19.77–19.99 | 0.00640 | -0.00033 | 0.01683 | 0 |

**Table S7**. CA intervals, mean AUC values and 95% CI bounds of the comparison of T1-based (pre-pandemic) normative developmental trajectory vs. T2 (post-pandemic) observed trajectory over the T2 age range. Significance of each CA window at a level of α<0.05 (two-sided) is indicated in the last column.

### Vulnerability scales with maturity-related momentum in the adolescent brain - Supplementary Tables S8-S14

| **Maturity group (T1)** | **N** | **CI-** | **CI+** | **IVWM** | **IVWV** | **CI length** | **CI (+, -)** | **SD** | **SE** | **BA-CA** |
| --- | --- | --- | --- | --- | --- | --- | --- | --- | --- | --- |
| Decelerated | 31 | 0.0158 | 0.0204 | 0.0181 | 0.0000014 | 0.0046 | 0.0023 | 0.0065 | 0.0012 | -1.01 |
| Average | 28 | 0.0213 | 0.0267 | 0.0240 | 0.0000019 | 0.0054 | 0.0027 | 0.0073 | 0.0014 | -0.01 |
| Accelerated | 28 | 0.0170 | 0.0219 | 0.0194 | 0.0000016 | 0.0049 | 0.0025 | 0.0066 | 0.0013 | 0.94 |

**Table S8.** Group-level descriptive statistics for T1 maturity groups, including number of participants, CI bounds, inverse-variance weighted mean (IVWM), inverse-variance weighted variance (IVWV), group-level standard deviation (SD), standard error (SE), and BA–CA (maturational timing).

| **Two-sample z-tests (T1)** | **Decelerated vs. average** | **Accelerated vs. average** | **Decelerated vs. accelerated** |
| --- | --- | --- | --- |
| z-value | 3.25 | 2.44 | 0.78 |
| p-value | **0.001** | **0.015** | 0.437 |
| Hedge's g | 0.85 | 0.65 | 0.20 |
|  | large | medium | small |

**Table S9.** Two-sample z-test results comparing T1 maturity groups, with corresponding Hedges’ g effect sizes. Results significant at α<0.05 are bold.

| **Maturity group (T2)** | **N** | **CI-** | **CI+** | **IVWM** | **IVWV** | **SD** | **SE** | **Age** |
| --- | --- | --- | --- | --- | --- | --- | --- | --- |
| Decelerated | 19 | 0.0163 | 0.0221 | 0.0192 | 0.0000022 | 0.0064 | 0.0015 | 18.53 |
| Average | 18 | 0.0186 | 0.0251 | 0.0219 | 0.0000028 | 0.0071 | 0.0017 | 18.06 |
| Accelerated | 19 | 0.0146 | 0.0201 | 0.0174 | 0.0000019 | 0.0060 | 0.0014 | 18.12 |

**Table S10.** Group-level descriptive statistics for T2 maturity groups (based on T1 assessment), including number of participants, CI bounds, IVWM, IVWV, group-level standard deviation (SD), standard error (SE), and mean CA.

| **Two-sample z-tests (T2)** | **Decelerated vs. average** | **Accelerated vs. average** | **Decelerated vs. accelerated** |
| --- | --- | --- | --- |
| z-value | 1.19 | 02.08 | 0.91 |
| p-value | 0.234 | **0.038** | 0.363 |
| Hedge's g | 0.39 | 0.69 | 0.30 |
|  | small | medium | small |

**Table S11.** Two-sample z-test results comparing maturity groups at T2, with corresponding Hedges’ g effect sizes. Results significant at α<0.05 are bold.

### Group comparisons using random-effects weighting and unweighted values – Fig. S1

**Fig. S1. Mean group-level entropy production with 95% confidence intervals within maturity groups at T1 and T2.**

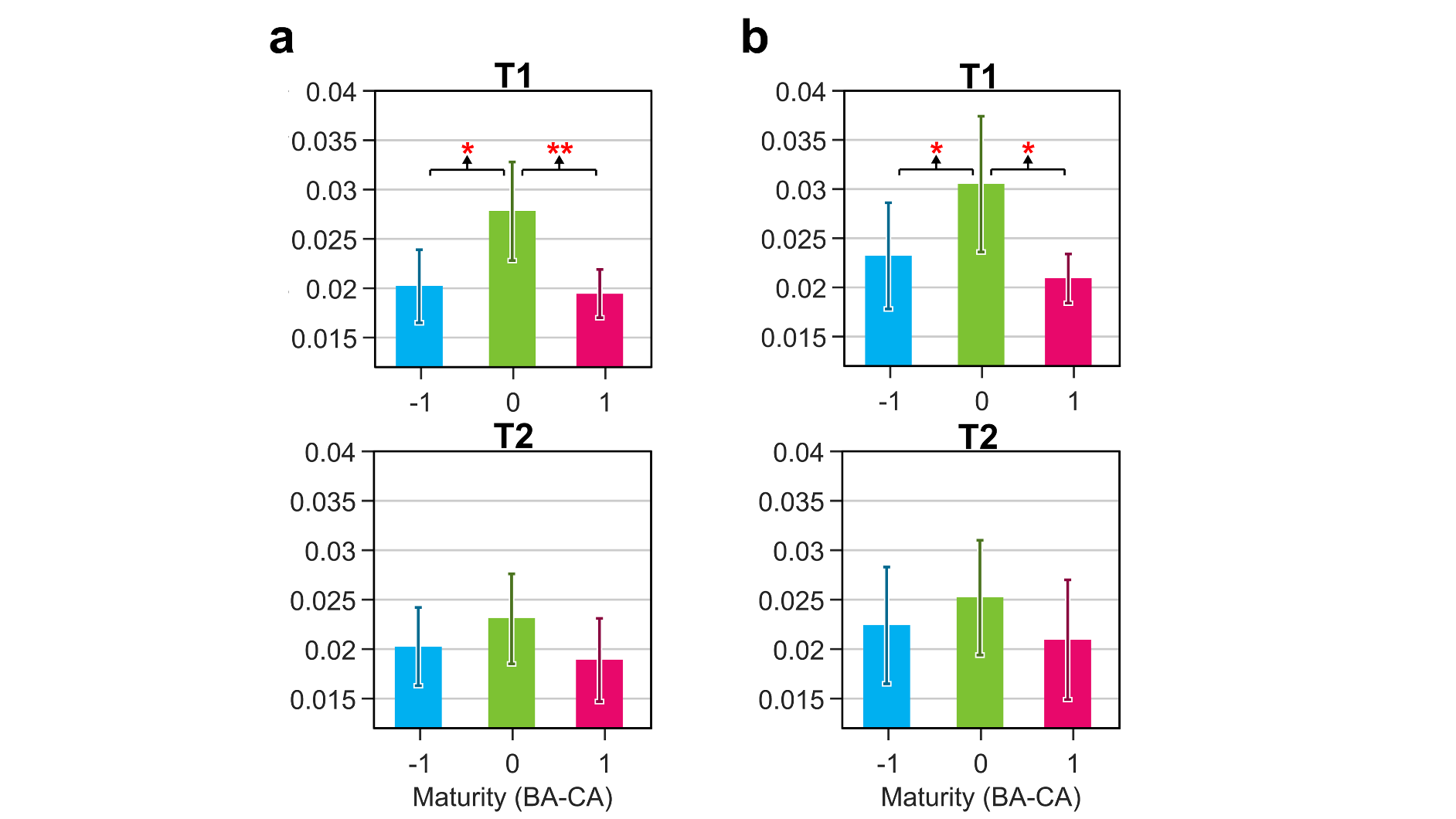

Means were estimated using random-effects weighting (column **a**), and unweighted values (column **b**). Pairwise comparisons (z-tests for random-effects weighted means; Mann–Whitney U tests for unweighted group comparisons) were tested at α = 0.05. **a**, Random-effects weighted means were calculated using the DerSimonian–Laird estimator of between-participant variance (τ²) computed separately for each group (DerSimonian & Laird, 1986). The average maturity group (0.0278 bit; 95% CI: 0.0228–0.0328) showed significantly higher group-level entropy production compared to both the decelerated group (0.0202 bit; 95% CI: 0.0165–0.0239; *p*=0.017; Hedges' *g*: 0.47, small effect) and the accelerated group (0.0194 bit; 95% CI: 0.0170–0.0219; *p*=0.003; Hedges' *g*: 0.63, medium effect) at T1. At T2, the comparison of the average (0.0231 bit; 95% CI: 0.0185–0.0276) and decelerated maturity groups (0.0202 bit; 95% CI: 0.0163–0.0242; *p*=0.362), and the comparison of the average and accelerated maturity groups (0.0189 bit; 95% CI: 0.0147–0.0231; *p*=0.188) did not yield significant results. **b**, Mann-Whitney U tests revealed significant differences between the average (0.0305 bit; 95% CI: 0.0236–0.0374) and decelerated (0.0232 bit; 95% CI: 0.0178–0.0286; *U*=279.000, *p*=0.019, r=0.31, medium effect) groups, and between the average and accelerated (0.0209 bit; 95% CI: 0.0184–0.0234; *U* =265.000, *p*=0.037, r=0.28, small effect) groups. At T2, the comparison of the average (0.0252 bit; 95% CI: 0.0194–0.0310) and decelerated maturity groups (0.0224 bit; 95% CI: 0.0165–0.0283; *p*=0.378), and the comparison of the average and accelerated maturity groups (0.0209 bit; 95% CI: 0.0149–0.0270; *p*=0.181) did not yield significant results.

| **CA interval** | **Mean AUC** | **95% CI (lower)** | **95% CI (upper)** | **Significant (1: yes; 0: no)** |
| --- | --- | --- | --- | --- |
| 16.91–17.16 | 0.00081 | -0.00116 | 0.00314 | 0 |
| 17.17–17.42 | 0.00089 | -0.00119 | 0.00341 | 0 |
| 17.43–17.68 | 0.00090 | -0.00114 | 0.00339 | 0 |
| 17.69–17.94 | 0.00098 | -0.00118 | 0.00370 | 0 |
| 17.95–18.20 | 0.00098 | -0.00113 | 0.00371 | 0 |
| 18.21–18.46 | 0.00107 | -0.00116 | 0.00405 | 0 |
| 18.47–18.72 | 0.00106 | -0.00110 | 0.00409 | 0 |
| 18.73–18.98 | 0.00115 | -0.00115 | 0.00446 | 0 |
| 18.99–19.24 | 0.00115 | -0.00110 | 0.00445 | 0 |
| 19.25–19.50 | 0.00119 | -0.00109 | 0.00461 | 0 |
| 19.51–19.76 | 0.00128 | -0.00113 | 0.00502 | 0 |
| 19.77–19.99 | 0.00111 | -0.00095 | 0.00439 | 0 |

**Table S12.** CA intervals, mean AUC values and 95% CI bounds of the comparison of T1-based (pre-pandemic) normative developmental trajectory vs. T2 (post-pandemic) observed trajectory of the accelerated maturity group over the T2 age range. Significance of each CA window at a level of α<0.05 (two-sided) is indicated in the last column.

| **CA interval** | **Mean AUC** | **95% CI (lower)** | **95% CI (upper)** | **Significant (1: yes; 0: no)** |
| --- | --- | --- | --- | --- |
| 16.91–17.16 | 0.00701 | 0.00051 | 0.01652 | 1 |
| 17.17–17.42 | 0.00743 | 0.00042 | 0.01804 | 1 |
| 17.43–17.68 | 0.00724 | 0.00025 | 0.01819 | 1 |
| 17.69–17.94 | 0.00765 | 0.00013 | 0.01975 | 1 |
| 17.95–18.20 | 0.00744 | -0.00010 | 0.01974 | 0 |
| 18.21–18.46 | 0.00786 | -0.00049 | 0.02143 | 0 |
| 18.47–18.72 | 0.00763 | -0.00100 | 0.02137 | 0 |
| 18.73–18.98 | 0.00805 | -0.00161 | 0.02314 | 0 |
| 18.99–19.24 | 0.00781 | -0.00208 | 0.02308 | 0 |
| 19.25–19.50 | 0.00789 | -0.00275 | 0.02387 | 0 |
| 19.51–19.76 | 0.00831 | -0.00350 | 0.02564 | 0 |
| 19.77–19.99 | 0.00704 | -0.00352 | 0.02219 | 0 |

**Table S13.** CA intervals, mean AUC values and 95% CI bounds of the comparison of T1-based (pre-pandemic) normative developmental trajectory vs. T2 (post-pandemic) observed trajectory of the average maturity group over the T2 age range. Significance of each CA window at a level of α<0.05 (two-sided) is indicated in the last column.

| **CA interval** | **Mean AUC** | **95% CI (lower)** | **95% CI (upper)** | **Significant (1: yes; 0: no)** |
| --- | --- | --- | --- | --- |
| 16.91–17.16 | 0.00960 | 0.00318 | 0.01793 | 1 |
| 17.17–17.42 | 0.01031 | 0.00310 | 0.01994 | 1 |
| 17.43–17.68 | 0.01017 | 0.00274 | 0.02031 | 1 |
| 17.69–17.94 | 0.01087 | 0.00257 | 0.02240 | 1 |
| 17.95–18.20 | 0.01068 | 0.00209 | 0.02263 | 1 |
| 18.21–18.46 | 0.01137 | 0.00186 | 0.02473 | 1 |
| 18.47–18.72 | 0.01113 | 0.00134 | 0.02472 | 1 |
| 18.73–18.98 | 0.01181 | 0.00103 | 0.02690 | 1 |
| 18.99–19.24 | 0.01152 | 0.00052 | 0.02692 | 1 |
| 19.25–19.50 | 0.01170 | 0.00004 | 0.02794 | 1 |
| 19.51–19.76 | 0.01236 | -0.00059 | 0.03020 | 0 |
| 19.77–19.99 | 0.01050 | -0.00097 | 0.02627 | 0 |

**Table S14.** CA intervals, mean AUC values and 95% CI bounds of the comparison of T1-based (pre-pandemic) normative developmental trajectory vs. T2 (post-pandemic) observed trajectory of the decelerated maturity group over the T2 age range. Significance of each CA window at a level of α<0.05 (two-sided) is indicated in the last column.

### Sensitivity analyses using 0.5, 1.0, and 1.5-year windows – Tables S15-S24

| **Windowing approach (years)** | **CA interval** | **Mean AUC** | **95% CI (lower)** | **95% CI (upper)** | **Significant (1: yes; 0: no)** |
| --- | --- | --- | --- | --- | --- |
| 0.50 | 11.60–12.10 | -0.00242 | -0.00802 | 0.00289 | 0 |
| 0.50 | 12.11–12.61 | -0.00311 | -0.00751 | 0.00114 | 0 |
| 0.50 | 12.62–13.12 | -0.00387 | -0.00757 | -0.00059 | 1 |
| 0.50 | 13.13–13.63 | -0.00463 | -0.00810 | -0.00138 | 1 |
| 0.50 | 13.64–14.14 | -0.00564 | -0.01012 | -0.00163 | 1 |
| 0.50 | 14.15–14.65 | -0.00663 | -0.01266 | -0.00153 | 1 |
| 0.50 | 14.66–15.16 | -0.00769 | -0.01588 | -0.00133 | 1 |
| 0.50 | 15.17–15.67 | -0.00882 | -0.01969 | -0.00103 | 1 |
| 0.50 | 15.68–15.97 | -0.00546 | -0.01272 | -0.00039 | 1 |
| 1.0 | 11.60–12.60 | -0.00551 | -0.01554 | 0.00408 | 0 |
| 1.0 | 12.61–13.61 | -0.00854 | -0.01553 | -0.00215 | 1 |
| 1.0 | 13.62–14.62 | -0.01216 | -0.02233 | -0.00318 | 1 |
| 1.0 | 14.63–15.97 | -0.02261 | -0.04960 | -0.00292 | 1 |
| 1.5 | 11.60–13.10 | -0.00936 | -0.02255 | 0.00328 | 0 |
| 1.5 | 13.11–14.61 | -0.01681 | -0.03013 | -0.00493 | 1 |
| 1.5 | 14.62–15.97 | -0.02289 | -0.05014 | -0.00297 | 1 |

**Table S15.** CA intervals, mean AUC values and 95% CI bounds for the comparison of the average vs. accelerated maturity group trajectories across the T1 age range, using 0.50-year, 1.0-year, and 1.5-year non-overlapping windows. Significance of each CA window at a level of α<0.05 (two-sided) is indicated in the last column.

| **Windowing approach (years)** | **CA interval** | **Mean AUC** | **95% CI (lower)** | **95% CI (upper)** | **Significant (1: yes; 0: no)** |
| --- | --- | --- | --- | --- | --- |
| 0.50 | 11.60–12.10 | 0.00587 | 0.00064 | 0.01098 | 1 |
| 0.50 | 12.11–12.61 | 0.00536 | 0.00085 | 0.00975 | 1 |
| 0.50 | 12.62–13.12 | 0.00475 | 0.00098 | 0.00864 | 1 |
| 0.50 | 13.13–13.63 | 0.00395 | 0.00038 | 0.00778 | 1 |
| 0.50 | 13.64–14.14 | 0.00326 | -0.00163 | 0.00824 | 0 |
| 0.50 | 14.15–14.65 | 0.00243 | -0.00439 | 0.00950 | 0 |
| 0.50 | 14.66–15.16 | 0.00155 | -0.00744 | 0.01133 | 0 |
| 0.50 | 15.17–15.67 | 0.00065 | -0.01123 | 0.01376 | 0 |
| 0.50 | 15.68–15.97 | -0.00003 | -0.00817 | 0.00882 | 0 |
| 1.0 | 11.60–12.60 | 0.01124 | 0.00147 | 0.02083 | 1 |
| 1.0 | 12.61–13.61 | 0.00883 | 0.00162 | 0.01647 | 1 |
| 1.0 | 13.62–14.62 | 0.00577 | -0.00546 | 0.01731 | 0 |
| 1.0 | 14.63–15.97 | 0.00228 | -0.02740 | 0.03486 | 0 |
| 1.5 | 11.60–13.10 | 0.01601 | 0.00269 | 0.02904 | 1 |
| 1.5 | 13.11–14.61 | 0.00988 | -0.00459 | 0.02461 | 0 |
| 1.5 | 14.62–15.97 | 0.00236 | -0.02766 | 0.03526 | 0 |

**Table S16**. CA intervals, mean AUC values and 95% CI bounds for the comparison of the average vs. decelerated maturity group trajectories across the T1 age range, using 0.50-year, 1.0-year, and 1.5-year non-overlapping windows. Significance of each CA window at a level of α<0.05 (two-sided) is indicated in the last column.

| **Windowing approach (years)** | **CA interval** | **Mean AUC** | **95% CI (lower)** | **95% CI (upper)** | **Significant (1: yes; 0: no)** |
| --- | --- | --- | --- | --- | --- |
| 0.50 | 11.60–12.10 | 0.00345 | 0.00003 | 0.00655 | 1 |
| 0.50 | 12.11–12.61 | 0.00225 | -0.00067 | 0.00512 | 0 |
| 0.50 | 12.62–13.12 | 0.00087 | -0.00182 | 0.00344 | 0 |
| 0.50 | 13.13–13.63 | -0.00068 | -0.00344 | 0.00185 | 0 |
| 0.50 | 13.64–14.14 | -0.00237 | -0.00577 | 0.00076 | 0 |
| 0.50 | 14.15–14.65 | -0.00420 | -0.00879 | 0.00019 | 0 |
| 0.50 | 14.66–15.16 | -0.00614 | -0.01215 | -0.00032 | 1 |
| 0.50 | 15.17–15.67 | -0.00817 | -0.01639 | -0.00069 | 1 |
| 0.50 | 15.68–15.97 | -0.00549 | -0.01114 | -0.00038 | 1 |
| 1.0 | 11.60–12.60 | 0.00572 | -0.00061 | 0.01172 | 0 |
| 1.0 | 12.61–13.61 | 0.00029 | -0.00515 | 0.00514 | 0 |
| 1.0 | 13.62–14.62 | -0.00639 | -0.01410 | 0.00097 | 0 |
| 1.0 | 14.63–15.97 | -0.02033 | -0.04071 | -0.00164 | 1 |
| 1.5 | 11.60–13.10 | 0.00665 | -0.00202 | 0.01518 | 0 |
| 1.5 | 13.11–14.61 | -0.00693 | -0.01688 | 0.00229 | 0 |
| 1.5 | 14.62–15.97 | -0.02053 | -0.04109 | -0.00165 | 1 |

**Table S17**. CA intervals, mean AUC values and 95% CI bounds for the comparison of the accelerated vs. decelerated maturity group trajectories across the T1 age range, using 0.50-year, 1.0-year, and 1.5-year non-overlapping windows. Significance of each CA window at a level of α<0.05 (two-sided) is indicated in the last column.

| **Windowing approach (years)** | **CA interval** | **Δslope** | **95% CI (lower)** | **95% CI (upper)** | **Significant (1: yes; 0: no)** |
| --- | --- | --- | --- | --- | --- |
| 0.50 | 11.60–12.10 | -0.00257 | -0.00750 | 0.00353 | 0 |
| 0.50 | 12.11–12.61 | -0.00285 | -0.00860 | 0.00327 | 0 |
| 0.50 | 12.62–13.12 | -0.00315 | -0.00994 | 0.00308 | 0 |
| 0.50 | 13.13–13.63 | -0.00346 | -0.01133 | 0.00285 | 0 |
| 0.50 | 13.64–14.14 | -0.00375 | -0.01259 | 0.00261 | 0 |
| 0.50 | 14.15–14.65 | -0.00403 | -0.01373 | 0.00252 | 0 |
| 0.50 | 14.66–15.16 | -0.00430 | -0.01471 | 0.00238 | 0 |
| 0.50 | 15.17–15.67 | -0.00455 | -0.01568 | 0.00228 | 0 |
| 0.50 | 15.68–15.97 | -0.00473 | -0.01630 | 0.00226 | 0 |
| 1.0 | 11.60–12.60 | -0.00271 | -0.00803 | 0.00343 | 0 |
| 1.0 | 12.61–13.61 | -0.00329 | -0.01050 | 0.00299 | 0 |
| 1.0 | 13.62–14.62 | -0.00387 | -0.01307 | 0.00259 | 0 |
| 1.0 | 14.63–15.97 | -0.00449 | -0.01540 | 0.00231 | 0 |
| 1.5 | 11.60–13.10 | -0.00285 | -0.00858 | 0.00324 | 0 |
| 1.5 | 13.11–14.61 | -0.00373 | -0.01248 | 0.00262 | 0 |
| 1.5 | 14.62–15.97 | -0.00449 | -0.01539 | 0.00231 | 0 |

**Table S18.** CA intervals, mean $\Delta$slope values, and 95% CI bounds for the comparison of the accelerated vs. average maturity groups at T1, using 0.50-year, 1.0-year, and 1.5-year non-overlapping windows. Significance of each CA window at a level of α<0.05 (two-sided) is indicated in the last column.

| **Windowing approach (years)** | **CA interval** | **Δslope** | **95% CI (lower)** | **95% CI (upper)** | **Significant (1: yes; 0: no)** |
| --- | --- | --- | --- | --- | --- |
| 0.50 | 11.60–12.10 | -0.00174 | -0.00674 | 0.00337 | 0 |
| 0.50 | 12.11–12.61 | -0.00220 | -0.00786 | 0.00404 | 0 |
| 0.50 | 12.62–13.12 | -0.00260 | -0.00923 | 0.00475 | 0 |
| 0.50 | 13.13–13.63 | -0.00293 | -0.01085 | 0.00577 | 0 |
| 0.50 | 13.64–14.14 | -0.00317 | -0.01247 | 0.00696 | 0 |
| 0.50 | 14.15–14.65 | -0.00336 | -0.01399 | 0.00785 | 0 |
| 0.50 | 14.66–15.16 | -0.00349 | -0.01564 | 0.00896 | 0 |
| 0.50 | 15.17–15.67 | -0.00356 | -0.01697 | 0.00998 | 0 |
| 0.50 | 15.68–15.97 | -0.00359 | -0.01803 | 0.01082 | 0 |
| 1.0 | 11.60–12.60 | -0.00197 | -0.00727 | 0.00364 | 0 |
| 1.0 | 12.61–13.61 | -0.00275 | -0.01000 | 0.00518 | 0 |
| 1.0 | 13.62–14.62 | -0.00326 | -0.01312 | 0.00739 | 0 |
| 1.0 | 14.63–15.97 | -0.00353 | -0.01661 | 0.00975 | 0 |
| 1.5 | 11.60–13.10 | -0.00217 | -0.00782 | 0.00403 | 0 |
| 1.5 | 13.11–14.61 | -0.00314 | -0.01228 | 0.00685 | 0 |
| 1.5 | 14.62–15.97 | -0.00353 | -0.01660 | 0.00974 | 0 |

**Table S19.** CA intervals, mean $\Delta$slope values, and 95% CI bounds for the comparison of the average vs. decelerated maturity groups at T1, using 0.50-year, 1.0-year, and 1.5-year non-overlapping windows. Significance of each CA window at a level of α<0.05 (two-sided) is indicated in the last column.

| **Windowing approach (years)** | **CA interval** | **Δslope** | **95% CI (lower)** | **95% CI (upper)** | **Significant (1: yes; 0: no)** |
| --- | --- | --- | --- | --- | --- |
| 0.50 | 11.60–12.10 | -0.00430 | -0.00748 | -0.00028 | 1 |
| 0.50 | 12.11–12.61 | -0.00506 | -0.00857 | -0.00071 | 1 |
| 0.50 | 12.62–13.12 | -0.00575 | -0.00975 | -0.00104 | 1 |
| 0.50 | 13.13–13.63 | -0.00638 | -0.01149 | -0.00114 | 1 |
| 0.50 | 13.64–14.14 | -0.00692 | -0.01325 | -0.00108 | 1 |
| 0.50 | 14.15–14.65 | -0.00739 | -0.01511 | -0.00104 | 1 |
| 0.50 | 14.66–15.16 | -0.00779 | -0.01700 | -0.00073 | 1 |
| 0.50 | 15.17–15.67 | -0.00811 | -0.01848 | -0.00049 | 1 |
| 0.50 | 15.68–15.97 | -0.00832 | -0.01962 | -0.00029 | 1 |
| 1.0 | 11.60–12.60 | -0.00467 | -0.00799 | -0.00049 | 1 |
| 1.0 | 12.61–13.61 | -0.00605 | -0.01054 | -0.00112 | 1 |
| 1.0 | 13.62–14.62 | -0.00713 | -0.01406 | -0.00109 | 1 |
| 1.0 | 14.63–15.97 | -0.00802 | -0.01806 | -0.00054 | 1 |
| 1.5 | 11.60–13.10 | -0.00502 | -0.00846 | -0.00072 | 1 |
| 1.5 | 13.11–14.61 | -0.00686 | -0.01306 | -0.00110 | 1 |
| 1.5 | 14.62–15.97 | -0.00802 | -0.01805 | -0.00054 | 1 |

**Table S20.** CA intervals, mean $\Delta$slope values, and 95% CI bounds for the comparison of the accelerated vs. decelerated maturity groups at T1, using 0.50-year, 1.0-year, and 1.5-year non-overlapping windows. Significance of each CA window at a level of α<0.05 (two-sided) is indicated in the last column.

| **Windowing approach (years)** | **CA interval** | **Mean AUC** | **95% CI (lower)** | **95% CI (upper)** | **Significant (1: yes; 0: no)** |
| --- | --- | --- | --- | --- | --- |
| 0.50 | 16.91–17.41 | 0.00968 | 0.00101 | 0.02215 | 1 |
| 0.50 | 17.42–17.92 | 0.01083 | 0.00085 | 0.02556 | 1 |
| 0.50 | 17.93–18.43 | 0.01156 | 0.00064 | 0.02841 | 1 |
| 0.50 | 18.44–18.94 | 0.01281 | 0.00029 | 0.03249 | 1 |
| 0.50 | 18.95–19.45 | 0.01355 | -0.00013 | 0.03480 | 0 |
| 0.50 | 19.46–19.99 | 0.01551 | -0.00064 | 0.04052 | 0 |
| 1.0 | 16.91–17.91 | 0.02048 | 0.00193 | 0.04745 | 1 |
| 1.0 | 17.92–18.92 | 0.02455 | 0.00081 | 0.06116 | 1 |
| 1.0 | 18.93–19.99 | 0.03016 | -0.00066 | 0.07788 | 0 |
| 1.5 | 16.91–18.41 | 0.03222 | 0.00256 | 0.07601 | 1 |
| 1.5 | 18.42–19.99 | 0.04319 | -0.00042 | 0.11085 | 0 |

**Table S21**. CA intervals, mean AUC values and 95% CI bounds of the comparison of T1-based (pre-pandemic) normative developmental trajectory vs. T2 (post-pandemic) observed trajectory over the T2 age range, using 0.50-year, 1.0-year, and 1.5-year non-overlapping windows. Significance of each CA window at a level of α<0.05 (two-sided) is indicated in the last column.

| **Windowing approach (years)** | **CA interval** | **Mean AUC** | **95% CI (lower)** | **95% CI (upper)** | **Significant (1: yes; 0: no)** |
| --- | --- | --- | --- | --- | --- |
| 0.50 | 16.91–17.41 | 0.00170 | -0.00233 | 0.00656 | 0 |
| 0.50 | 17.42–17.92 | 0.00191 | -0.00236 | 0.00718 | 0 |
| 0.50 | 17.93–18.43 | 0.00204 | -0.00230 | 0.00772 | 0 |
| 0.50 | 18.44–18.94 | 0.00225 | -0.00230 | 0.00871 | 0 |
| 0.50 | 18.95–19.45 | 0.00237 | -0.00224 | 0.00917 | 0 |
| 0.50 | 19.46–19.99 | 0.00269 | -0.00236 | 0.01060 | 0 |
| 1.0 | 16.91–17.91 | 0.00360 | -0.00471 | 0.01378 | 0 |
| 1.0 | 17.92–18.92 | 0.00432 | -0.00463 | 0.01649 | 0 |
| 1.0 | 18.93–19.99 | 0.00526 | -0.00477 | 0.02045 | 0 |
| 1.5 | 16.91–18.41 | 0.00567 | -0.00704 | 0.02133 | 0 |
| 1.5 | 18.42–19.99 | 0.00755 | -0.00712 | 0.02923 | 0 |

**Table S22.** CA intervals, mean AUC values and 95% CI bounds of the comparison of T1-based (pre-pandemic) normative developmental trajectory vs. T2 (post-pandemic) observed trajectory of the accelerated maturity group over the T2 age range, using 0.50-year, 1.0-year, and 1.5-year non-overlapping windows. Significance of each CA window at a level of α<0.05 (two-sided) is indicated in the last column.

| **Windowing approach (years)** | **CA interval** | **Mean AUC** | **95% CI (lower)** | **95% CI (upper)** | **Significant (1: yes; 0: no)** |
| --- | --- | --- | --- | --- | --- |
| 0.50 | 16.91–17.41 | 0.01443 | 0.00095 | 0.03458 | 1 |
| 0.50 | 17.42–17.92 | 0.01518 | 0.00042 | 0.03860 | 1 |
| 0.50 | 17.93–18.43 | 0.01528 | -0.00052 | 0.04087 | 0 |
| 0.50 | 18.44–18.94 | 0.01598 | -0.00256 | 0.04522 | 0 |
| 0.50 | 18.95–19.45 | 0.01600 | -0.00480 | 0.04774 | 0 |
| 0.50 | 19.46–19.99 | 0.01734 | -0.00770 | 0.05382 | 0 |
| 1.0 | 16.91–17.91 | 0.02960 | 0.00144 | 0.07300 | 1 |
| 1.0 | 17.92–18.92 | 0.03155 | -0.00272 | 0.08702 | 0 |
| 1.0 | 18.93–19.99 | 0.03464 | -0.01285 | 0.10549 | 0 |
| 1.5 | 16.91–18.41 | 0.04517 | 0.00127 | 0.11473 | 1 |
| 1.5 | 18.42–19.99 | 0.05093 | -0.01543 | 0.15195 | 0 |

**Table S23.** CA intervals, mean AUC values and 95% CI bounds of the comparison of T1-based (pre-pandemic) normative developmental trajectory vs. T2 (post-pandemic) observed trajectory of the average maturity group over the T2 age range, using 0.50-year, 1.0-year, and 1.5-year non-overlapping windows. Significance of each CA window at a level of α<0.05 (two-sided) is indicated in the last column.

| **Windowing approach (years)** | **CA interval** | **Mean AUC** | **95% CI (lower)** | **95% CI (upper)** | **Significant (1: yes; 0: no)** |
| --- | --- | --- | --- | --- | --- |
| 0.50 | 16.91–17.41 | 0.01989 | 0.00624 | 0.03765 | 1 |
| 0.50 | 17.42–17.92 | 0.02144 | 0.00544 | 0.04346 | 1 |
| 0.50 | 17.93–18.43 | 0.02199 | 0.00409 | 0.04715 | 1 |
| 0.50 | 18.44–18.94 | 0.02335 | 0.00250 | 0.05231 | 1 |
| 0.50 | 18.95–19.45 | 0.02364 | 0.00073 | 0.05559 | 1 |
| 0.50 | 19.46–19.99 | 0.02581 | -0.00160 | 0.06345 | 0 |
| 1.0 | 16.91–17.91 | 0.04128 | 0.01172 | 0.08098 | 1 |
| 1.0 | 17.92–18.92 | 0.04575 | 0.00687 | 0.10018 | 1 |
| 1.0 | 18.93–19.99 | 0.05137 | -0.00071 | 0.12376 | 0 |
| 1.5 | 16.91–18.41 | 0.06368 | 0.01609 | 0.12905 | 1 |
| 1.5 | 18.42–19.99 | 0.07516 | 0.00217 | 0.17687 | 1 |

**Table S24.** CA intervals, mean AUC values and 95% CI bounds of the comparison of T1-based (pre-pandemic) normative developmental trajectory vs. T2 (post-pandemic) observed trajectory of the decelerated maturity group over the T2 age range, using 0.50-year, 1.0-year, and 1.5-year non-overlapping windows. Significance of each CA window at a level of α<0.05 (two-sided) is indicated in the last column.

### Additional demographic data - Tables S25-S27

| **Parental education** |  |  |  |  |  |  |  |
| --- | --- | --- | --- | --- | --- | --- | --- |
|  | Mother |  | **Less than 8 grades** | **8 grades** | **Vocational school** | **High school** | **College or university** |
|  |  | Education | 1 | 2 | 3 | 4 | 5 |
|  |  | N | 0 | 0 | 0 | 5 | 81 |
|  |  | Percentage | 0% | 0% | 0% | 6% | 93% |
|  | Father |  | **Less than 8 grades** | **8 grades** | **Vocational school** | **High school** | **College or university** |
|  |  | Education | 1 | 2 | 3 | 4 | 5 |
|  |  | N | 0 | 0 | 0 | 6 | 80 |
|  |  | Percentage | 0% | 0% | 0% | 7% | 92% |

**Table S25.** Parental education of participants, measured at T1.

| **Place of residence (T1)** |  | **Capital city** | **Large city** | **Small town** | **Village** |
| --- | --- | --- | --- | --- | --- |
|  | Residence type (code) | 1 | 2 | 3 | 4 |
|  | N | 73 | 3 | 5 | 2 |
|  | Percentage | 84% | 3% | 6% | 2% |

| **School type (T1)** |  | **Standard 8-grade primary school** | **Alternative curriculum 8-grade primary school** | **6-grade high school** | **8-grade high school** | **Other** |
| --- | --- | --- | --- | --- | --- | --- |
|  | School type (code) | 1 | 2 | 3 | 4 | 5 |
|  | N | 1 | 0 | 27 | 52 | 0 |
|  | Percentage | 1% | 0% | 31% | 60% | 0% |

**Table S26.** Place of residence and school type of participants at T1.

|  |  |  | **Yes** | **No** | **⅀** | **Percentage of post-menarche participants** |
| --- | --- | --- | --- | --- | --- | --- |
| **Menarche per maturity group (T1)** | | Decelerated | 15 | 16 | 31 | 48% |
|  |  | Average | 21 | 7 | 28 | 75% |
|  |  | Accelerated | 21 | 7 | 28 | 75% |

**Table S27.** Percentage of post-menarche participants per maturity group at T1.

### Data on COVID-19 infection and perceived effects on life domains – Table S28

| **COVID-19 infection and vaccination history (N = 36)** | | | |
| --- | --- | --- | --- |
|  | **N** | **%** |  |
| Personal COVID-19 infection | 15 | 41.7 |  |
| COVID-19 infection in the family | 32 | 88.9 |  |
| COVID-19 vaccination | 35 | 97.2 |  |
| **Post-COVID symptoms among infected participants (N = 14)** | | | |
|  | **N** | **%** |  |
| Symptoms persisting ≥ 6 months | 11 | 78.6 |  |
| No persisting symptoms | 3 | 21.4 |  |
| **PCSS-C/Y symptom categories among infected participants (N = 14)** | | | |
| **Symptom category (no. of items)** | **0 symptoms (%)** | **≥ 1 symptom (%)** | **Mean (SD)** |
| Neurocognitive (4) | 78.6 | 21.4 | 0.57 (1.22) |
| Cardiorespiratory (4) | 64.3 | 35.7 | 0.64 (1.01) |
| Olfactory (2) | 78.6 | 21.4 | 0.36 (0.74) |
| Non-specific somatic (7) | 35.7 | 64.3 | 1.14 (1.29) |
| **Self-reported effects of pandemic measures on life domains (N = 36)** | | | |
| **Domain** | **Adverse (%)** | **Neutral (%)** | **Favourable (%)** |
| Life as a whole | 52.8 | 33.3 | 13.9 |
| Health | 22.2 | 61.1 | 16.7 |
| Relationships with family | 36.1 | 30.6 | 33.3 |
| Relationships with friend | 66.7 | 25.0 | 8.3 |
| Mental health | 61.1 | 19.4 | 19.4 |
| School performance | 36.1 | 38.9 | 25.0 |
| Physical activity | 47.2 | 16.7 | 36.1 |
| Eating and drinking | 38.9 | 38.9 | 22.2 |
| Future expectations* | 25.7 | 62.9 | 11.4 |
| Family’s financial situation | 19.4 | 72.2 | 8.3 |

**Table S28.** Summary of retrospective survey responses on pandemic-related experiences in the T2 sample (N = 36, 64% response rate). The survey was conducted between May and October 2025 and included items on COVID-19 infection and vaccination history, the Post-COVID Symptom Scale for Children/Youth^1^ (PCSS-C/Y), as well as items from the Health Behaviour in School-aged Children (HBSC^2^; Hungarian translation^3^) study assessing health and well-being during the pandemic. Responses were categorised following the approach of the 2022 HBSC study in Hungary: adverse (very unfavourable or quite unfavourable), neutral (neither favourable nor unfavourable), and favourable (quite favourable or very favourable). *N=35 (1 missing). The Hungarian United Ethical Review Committee for Research in Psychology (EPKEB) approved data collection using the listed questionnaires (reference number 2025-053).

### Assessment of selective follow-up – Table S29

| **Mann-Whitney U tests** | | | | |
| --- | --- | --- | --- | --- |
| **Baseline variable** | **Total N** | **Mean rank** | **Mann-Whitney U** | **P-value (2-tailed)** |
| Mean individual entropy production | 87 | Retained: 44.27 (N=56)  Not retained: 43.52 (N=31) | 883.000 | .894 |
| Chronological Age | 87 | Retained: 41.75 (N=56)  Not retained: 48.06 (N=31) | 742.000 | .264 |
| Maturity (BA-CA) | 87 | Retained: 45.25 (N=56)  Not retained: 41.74 (N=31) | 938.000 | .535 |
| EEG length | 87 | Retained: 49.27 (N=56)  Not retained: 34.48 (N=31) | 1163.000 | **.009** |
| Removed EEG channels | 87 | Retained: 44.73 (N=56)  Not retained: 42.68 (N=31) | 909.000 | .715 |
| Age at menarche | 57 | Retained: 26.63 (N=36)  Not retained: 33.07 (N=21) | 292.500 | .141 |
| Education of father | 86 | Retained: 43.37 (N=55)  Not retained: 43.73 (N=31) | 845.500 | .887 |
| Education of mother | 86 | Retained: 42.87 (N=55)  Not retained: 44.61 (N=31) | 818.000 | .444 |
| **χ² tests** | | | | |
| **Baseline variable** | **Total N** | **χ²** | **df** | **P-value (2-tailed)** |
| Place of residence | 83 | .177 | 3 | .981 |
| School type | 81 | 2.401 | 3 | .494 |
| Maturity group | 87 | .277 | 2 | .871 |

**Table S29.** Selective follow-up was assessed by comparing the listed variables, collected at T1, between retained (data acquisition at both timepoints) and non-retained (data acquisition at T1 only) participants. Retained participants did not significantly differ from those who did not return for follow-up in terms of mean individual entropy production values, chronological age, maturity (evaluated as BA-CA), number of removed EEG channels, age at menarche, parental education, place of residence, school type, or maturity group assignment. However, participants who returned for T2 measurements had significantly longer baseline EEG recordings following noisy epoch removal (Mann–Whitney U =1163, p=0.009). Overall, the T2 sample is representative of the T1 cohort.

### Distribution of removed EEG channels – Table S30

| **Sample** | **Group** | **Min** | **Max** | **Mean** | **SD** |
| --- | --- | --- | --- | --- | --- |
| T1 | Full T1 | 0 | 15 | 4.79 | 3.34 |
|  | Accelerated | 0 | 15 | 5.04 | 3.87 |
|  | Average | 0 | 10 | 4.75 | 2.84 |
|  | Decelerated | 0 | 13 | 4.61 | 3.33 |
| T2 | Full T2 | 0 | 8 | 2.30 | 2.46 |
|  | Accelerated | 0 | 8 | 2.74 | 2.73 |
|  | Average | 0 | 7 | 2.28 | 2.27 |
|  | Decelerated | 0 | 8 | 1.89 | 2.42 |

**Table S30**. Distribution of removed EEG channels across maturity groups at T1 and T2. Total number of electrodes = 128. Removed channels represent 0–12% of the total. According to Kruskal-Wallis rank tests, maturity groups did not differ in terms of removed channels (T1: *H(2)*=0.218, *p*=0.8966; T2: *H*(2)=1.085, *p*=0.5814).

### Comparison of Gompertz and linear model fits – Table S31 and Fig. S2

| **Subsample** | **N** | ***D*** | ***p*** |
| --- | --- | --- | --- |
| Full T1 | 87 | 0.0255 | 0.529 |
| Accelerated | 28 | 0.0225 | 0.687 |
| Average | 28 | 0.0255 | 0.529 |
| Decelerated | 31 | 0.0225 | 0.687 |

**Table S31.** Formal sensitivity analysis of Gompertz and linear fits. The table shows the results of two-sample Kolmogorov–Smirnov tests, comparing the distributions of the sum of squared residuals (SSR) obtained from Gompertz and linear functions fitted to the same 2000 bootstrap resamples within each subsample (Full T1; accelerated; average; decelerated). Rows correspond to panels a–d of Figure S1, which shows these distributions expressed as RMSE=√(SSR/N).

**Fig. S2. Comparison of Gompertz and linear fits across 2000 bootstrap resamples of the T1 sample.**

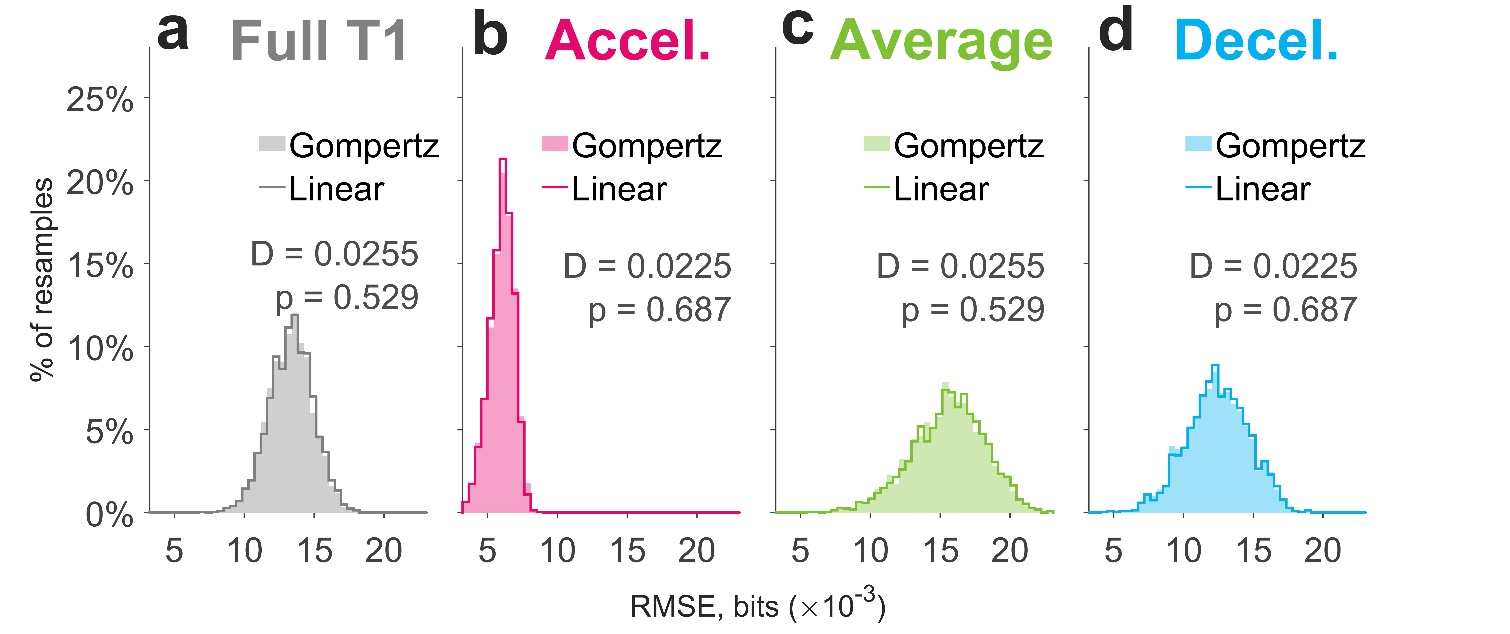

Each panel shows the distribution of goodness-of-fit values obtained from the Gompertz function (filled bars) and from a linear function (step outline) fitted to the same resampled data: **a**, the full T1 sample (grey), **b**, the accelerated (pink), **c**, the average (green), and **d**, the decelerated (blue) maturity group. Fit is normalized as RMSE (bits; x-axis)=√(SSR/N), a strictly monotone transformation, where SSR is the sum of squared residuals and N is the subsample size, placing subsamples of differing size on comparable axes. Bin edges are identical across all panels, and the y-axis shows the percentage of the 2000 resamples falling in each bin. Statistics derived from Kolmogorov–Smirnov tests, computed on the untransformed SSR distributions, are shown in each panel.

2.	HBSC study | Health Behaviour in School-aged Children study. https://www.hbsc.org/.

3.	*Iskoláskorú gyermekek egészségmagatartása 2022 : A WHO-val együttműködésben megvalósuló nemzetközi kutatás nemzeti jelentése – legújabb adatok és trendek az elmúlt húsz évben*. vol. 23 (L’Harmattan Kiadó, ELTE Pedagógiai és Pszichológiai Kar, 2024).
